# *Toxoplasma* IMC1 is a central component of the subpellicular network and plays critical roles in parasite morphology, replication, and infectivity

**DOI:** 10.1101/2025.06.03.657516

**Authors:** Juliette N. Uy, Qing Lou, Z. Hong Zhou, Peter J. Bradley

**Affiliations:** Department of Microbiology, Immunology, and Molecular Genetics, University of California, Los Angeles, Los Angeles, CA 90095; Department of Materials Science and Engineering, University of California, Los Angeles, Los Angeles, CA 90095; Molecular Biology Institute, University of California, Los Angeles, Los Angeles, CA 90095

**Keywords:** Toxoplasma gondii, Plasmodium, alveolin, inner membrane complex, IMC1, IMC4

## Abstract

*Toxoplasma gondii* and related apicomplexan parasites utilize a unique membrane and cytoskeletal organelle called the inner membrane complex (IMC) for maintaining cell shape, motility, host cell invasion, and replication. The cytoskeleton portion of the organelle is a network of filaments composed of proteins called alveolins, whose precise functions and organization are poorly understood. Here we describe the function of the founding member of the *Toxoplasma* alveolins, IMC1, which we show is expressed and loaded onto forming daughter buds with IMC4, but later than the other key alveolins IMC3, IMC6, and IMC10. Disruption of IMC1 results in severe morphological defects that impact the integrity of the parasite’s cytoskeleton and disrupt invasion, replication, and egress. Loss of IMC1 in a less virulent type II strain results in a dramatic loss of infectivity and complete failure to form a chronic infection. We then use deletion analyses to dissect functional regions of the protein which reveals a key subregion of the alveolin domain that is sufficient for IMC targeting and also required for function. We then show that IMC1 interacts directly with IMC4 and the loss of IMC1 results in mislocalization of IMC4 specifically in forming daughter buds. This study thus reveals the critical role that IMC1 plays in forming and maintaining the architecture of the filamentous network of the IMC.

**Significance:** Parasites in the phylum Apicomplexa maintain their intracellular lifestyle using specialized organelles that mediate the lytic cycle of host cell invasion, intracellular replication, and egress. One of these organelles is the inner membrane complex (IMC), which consists of membrane vesicles supported by a cytoskeletal meshwork formed from proteins called alveolins. This study focuses on the first identified alveolin IMC1 and determines its precise function via expression timing, gene knockout, deletion and mutagenesis, partner identification, and in vivo infection studies. We show that this protein is critical to the ultrastructure of the parasite which is important for every stage of its lytic cycle. We also identify key regions of the protein that are important for localization, function, and interaction with another key alveolin, IMC4.

## Introduction

The phylum Apicomplexa is composed of a diverse group of obligate intracellular parasites that are responsible for significant human and veterinary diseases worldwide (1). This includes the human pathogens *Toxoplasma gondii, Cryptosporidium spp*. and *Plasmodium spp.*, which cause toxoplasmosis, cryptosporidiosis, and malaria, respectively (2–4). Notable veterinary apicomplexans include *Neospora spp., Eimeria spp.,* and *Theileria spp.*, which cause livestock diseases that result in substantial economic losses worldwide (5–7). A defining feature of these parasites is the inner membrane complex (IMC), a specialized organelle that resides directly beneath the plasma membrane and plays critical roles in cell shape, motility, host cell invasion, and daughter cell formation during replication (8). Structurally, the IMC consists of a series of interconnected flattened alveolar vesicles supported by a cytoskeletal network known as the subpellicular network (SPN), which is a lattice of ∼8-10nm intermediate filaments composed of a family of alveolin-domain containing proteins (9). The IMC membranes and SPN are further supported by 22 underlying stable microtubules that emanate from the microtubule-based conoid at the apical end of the parasite (10).

The IMC is organized into several subdomains that contain distinct groups of proteins which carry out the diverse functions of the organelle. The alveolar membranes host the glideosome, an actin-myosin motor that bridges the IMC to the parasite’s plasma membrane via adhesins that are secreted onto the parasite’s surface from the micronemes (11). Action of the actin-myosin motor on the adhesins provides the driving force for active host cell invasion. The IMC also functions in daughter cell formation during the internal budding process of endodyogeny (12). The IMC membrane and SPN proteins are expressed in a “just in time” fashion in which proteins are sequentially expressed and added to the growing daughter buds within the cytoplasm of the maternal cell (13). This process is initiated by a trio of essential early daughter bud proteins (IMC32, IMC43, and BCC0) that form a complex named the “essential daughter bud assembly complex” (14–16). This complex serves as a foundation for the addition of sequentially expressed proteins to both the IMC membranes and the forming SPN, which enable maturation of the daughter cells. Another subdomain of the IMC with a unique function is the cone-shaped apical cap, which hosts proteins that are important for the regulation of the disassembly of the parasite’s cytoskeleton at the latest stages of replication (17–19). Finally, the posterior end of the IMC contains the basal complex, which plays critical roles in parasite replication and cytokinesis (20). While many of the proteins involved in these processes have been identified and characterized, their precise interactions and organization within the forming IMC and how these proteins are regulated remains largely unknown.

The SPN filaments are believed to be formed from a family of proteins called the alveolins, plus an assortment of alveolin-associated proteins (21). There are fourteen alveolins in *T. gondii* that range from a predicted 15.5-83.3 kDa, although the proteins typically migrate slower on SDS-PAGE gels than their calculated mass (22). The localizations of all fourteen alveolins have been determined, which shows several different localizations including the maternal IMC, maternal and daughter IMC, apical cap, and basal complex (23, 24). Each of the alveolins contain a conserved region of proline-valine rich repeats named the “alveolin domain” which frequently has ‘EKIVEVP’ and ‘EVVR’ or ‘VPV’ motifs (21). Deletion analyses of IMC3, IMC6 and IMC8 suggest the alveolin domain is largely sufficient for IMC targeting, but the N and C-terminal flanking regions also play a minor role in localization (23, 25). Many of the alveolins also have predicted palmitoylation sites in these regions, suggesting these members of the family are anchored in the SPN via their alveolin domain and then tethered to the IMC membranes via palmitoylation (23). The organization of the alveolins in the IMC is beginning to be better understood using unnatural amino acid (UAA) photocrosslinking, which can identify precise partners and binding interfaces, particularly in cellular environments that can be challenging for interaction studies such as the cytoskeleton (22, 25). These studies have shown that the IMC6 alveolin domain interacts with IMC3 and the essential non-alveolin protein ILP1 and that the ILP1 coiled coil domain interacts with IMC3, IMC6 and IMC27. Together, this suggests that the SPN components are highly interconnected with the alveolin domain playing a central role in filament assembly.

Data from the initial *T. gondii* genome-wide CRISPR screen indicates that IMC1, IMC3, IMC4, IMC6 and IMC10 are either essential or important for fitness, while the others are more likely to be dispensable (26). Intriguingly, these alveolins are present in the body of the maternal IMC and also in forming daughter buds, suggesting they are core members of the family that are critical for the construction of daughter buds and maintenance of the IMC ultrastructure (24). This is supported by our recent studies of *IMC6*, which can be disrupted, but the knockout results in substantial morphological and replication defects, resulting in diminished growth in vitro and dramatically reduced virulence in mice (25). Invasion is also impacted in Δ*imc6* parasites, but not severely, and egress is not, suggesting that the invasion defect may be linked to the rounding of extracellular parasites. IMC10 has also been investigated and shown to be important for tethering of the mitochondria to the IMC via interaction with the outer mitochondrial membrane protein LMF1 (27). Knockdown of IMC10 results in a loss of mitochondrial tethering with little other effects, but the lack of a bona fide IMC10 knockout suggests that other roles might still be present that can be supported by very low amounts of the protein. Other alveolins whose roles have been determined include IMC14, which functions in synchronous division, and IMC15, which plays a role in controlling the appropriate number of daughter buds during endodyogeny (24).

The first alveolin protein in *T. gondii* to be identified and initially characterized was IMC1 (28). IMC1 has a centrally located alveolin domain which is flanked by C and N-terminal regions of the protein that contain several predicted palmitoylation sites. The protein undergoes C-terminal proteolytic processing in the late daughter stages of parasite maturation, an event which has been reported to be important for firm attachment to the cytoskeleton (29). IMC1 has a low phenotype score in a genome-wide CRISPR screen, indicating that it plays a critical role in parasite fitness (26). In this paper, we directly assess the function of IMC1 and dissect regions of the protein that are sufficient for IMC targeting and function. We demonstrate that *IMC1* is surprisingly dispensable, but loss of the protein results in severe morphological defects that impact invasion, replication, and egress. Deletion analyses reveal that IMC1 only requires a small portion of the alveolin domain to properly localize to the IMC, and that this region is necessary for function. We also disrupted *IMC1* in type II strain parasites and demonstrate that the protein is essential for infectivity in vivo. We additionally show that IMC1 interacts directly with IMC4 and that loss of IMC1 results in a more fragile SPN. Together, this work highlights the critical role of IMC1 in partnering with IMC4 to stabilize parasite morphology and the cytoskeletal network of the SPN.

## Results

### IMC1 is recruited onto developing daughter buds concurrently with IMC4 and later than IMC3, 6, and 10

Among the fourteen alveolins that comprise the SPN, IMC1, IMC3, IMC4, IMC6, and IMC10 have been shown to be present in the IMC body of both maternal and budding parasites (28, 23, 24). While IMC1 and IMC4 are expressed equally in both the maternal and daughter cells, IMC3, IMC6, and IMC10 are enriched in the daughter cells during bud formation. Despite their localizations being previously identified, the timing of the recruitment of these proteins to the daughter bud cytoskeleton has yet to be characterized. To explore the precise timing of the expression and loading onto daughter buds of these key alveolins, we co-stained wild-type parasites with IMC1 and IMC3, IMC4, IMC6, and IMC10. We determined that IMC1 and IMC4 are recruited onto the growing daughter buds at equivalent times and always have overlapping localization patterns (Fig 1A). In contrast, we found that IMC3 (Fig 1B), IMC6 (Fig 1C), and IMC10 (Fig 1D) are all detected in daughter buds prior to the appearance of IMC1, suggesting that IMC1 and IMC4 are recruited to the growing IMC network later in replication than IMC3, IMC6, and IMC10. We also observed that IMC1 and IMC4 extend into the apical cap region of the IMC, whereas IMC3, IMC6 and IMC10 are restricted to the body portion of the IMC and are not present in the apical cap (Fig 1B, C, D).

**Figure 1.**
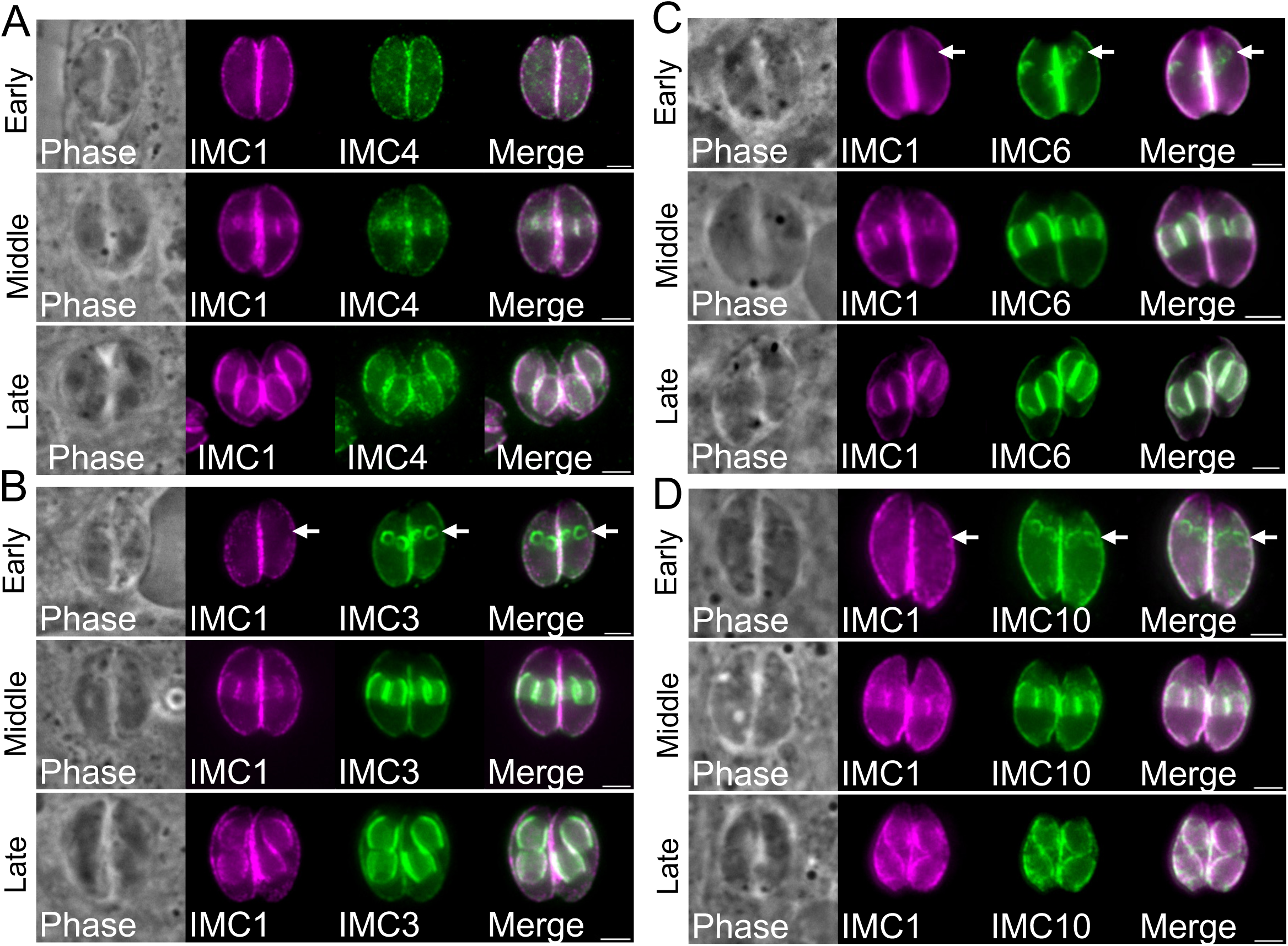
IMC1 is expressed later than IMC3, IMC6, and IMC10 and concurrently with IMC4. A) IFA showing IMC1 is expressed similar to IMC4 in early, middle, and late stages of daughter bud formation with mouse anti-IMC1 in magenta and rat anti-IMC4 in green. B) IFA showing IMC3 is expressed earlier than IMC1 during daughter bud formation. Arrows highlight the earlier expression in the early daughter buds panel. Mouse anti-IMC1 is in magenta and rabbit anti-IMC3 is in green. C) IFA showing IMC6 is expressed earlier than IMC1 during daughter bud formation stained with mouse anti-IMC1 in magenta and rabbit anti-IMC6 in green. D) IFA showing IMC1 is expressed earlier than IMC10 during daughter bud formation stained with mouse anti-IMC1 in magenta and rat anti-IMC10 in green. All scale bars are 2 µm. IFA, immunofluorescence assay.

### Disruption of IMC1 results in severe morphological defects *in vitro*

In the original *T.* gondii genome-wide CRISPR screen, *IMC1* was assigned a phenotype score of –4.0 (26), placing it among the lowest phenotype scores of the 14 alveolins. This highly negative score suggests essentiality; however, each gene must be individually assessed to determine whether it is truly essential. Thus, we attempted to disrupt *IMC1* in RHΔ*ku80*Δ*hxgprt* parasites and were surprisingly able to generate a line that lacked IMC1 staining by immunofluorescence assays, which also showed that the intracellular parasites appeared rounded and swollen (Δ*imc1*, Fig. 2A). The *IMC1* knockout was confirmed by PCR which showed an absence of the coding region and the presence of the selectable marker in the IMC1 locus in the knockout (Fig. 2B). We then rescued the loss of expression by inserting a full-length *IMC1* complementation construct driven by its endogenous promoter into the *UPRT* locus of the parasite (Fig. 2C) (30). We determined that the complemented protein localized similar to wild-type parasites by IFA (strain denoted IMC1^c^, Fig. 2D) and was also expressed at similar levels compared to the parental strain by western blot analysis (Fig. 2E).

**Figure 2.**
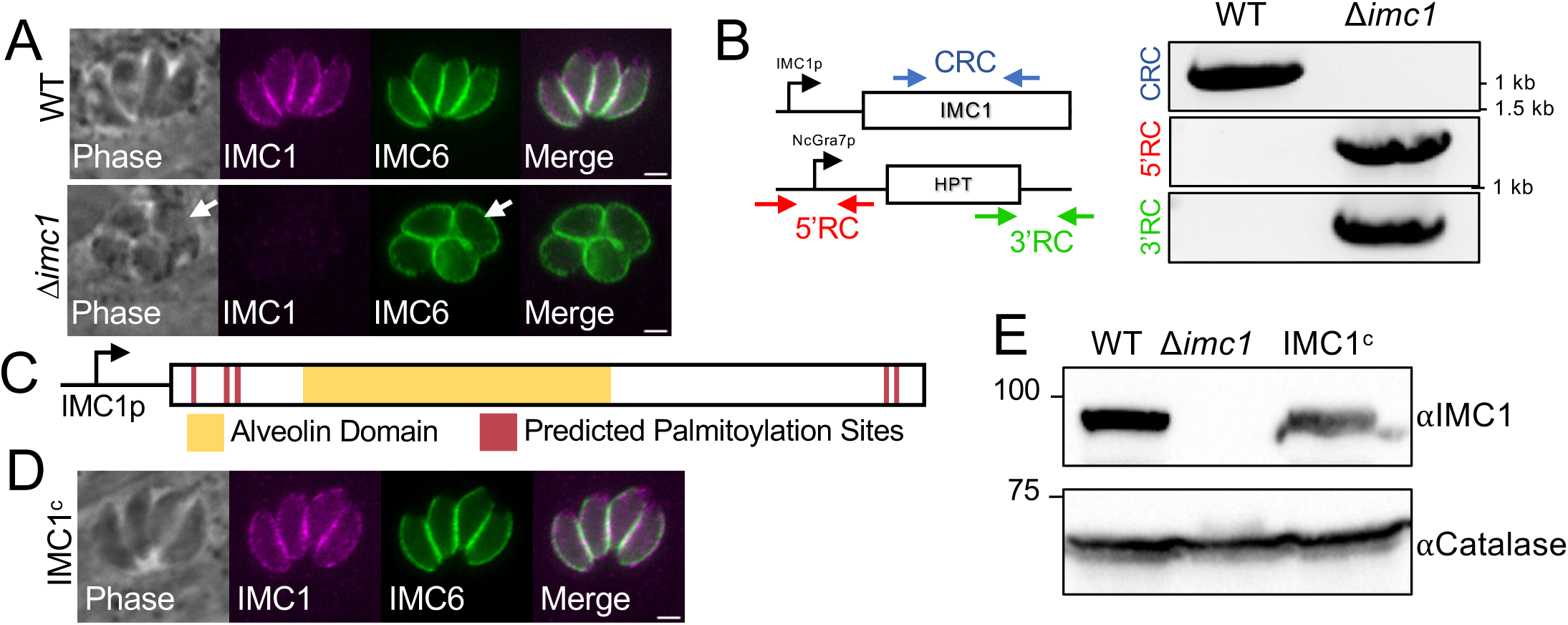
IMC1 knockout and complementation, loss of IMC1 affects parasite morphology. A) IFA of intracellular WT parasites showing proper localization of IMC1 (top). IFA of intracellular Δ*imc1* parasites showing the absence of IMC1 and swollen morphology (bottom, arrows). B) PCR verification of the genomic loci from WT (RhΔ*hxgprtΔku80*) and Δ*imc1* parasites. Diagram demonstrates primers used to amplify regions of the IMC1 coding sequence (blue arrows, coding region check) and the regions containing the 5’ and 3’ sites of recombination for the knockout locus (recombination check, red and green arrows). C) Schematic of the full-length IMC1 complementation construct driven by the endogenous IMC1 promoter. The full-length protein is 609 amino acids, with residues 98-422 (yellow) encompassing the alveolin domain. The five predicted palmitoylation sites with a GSS-Palm 4.0 cutoff score > 5 are shown (maroon). D) IFA of intracellular Δ*imc1* parasites expressing the IMC1 complementation construct, demonstrating restoration of IMC1 localization (IMC1^c^ strain). E) Western blots of whole cell lysates confirming similar protein levels expressed by the IMC1^c^ strain, as well as an absence of signal in the Δ*imc1* strain. Stained with mouse anti-IMC1 with rabbit anti-catalase serving as a load control. All scale bars are 2 µm. IFA, immunofluorescence assay; WT, wild-type; CRC, Coding Region Check; 5’RC, 5’ Recombination Check; 3’RC, 3’ Recombination Check.

### Disruption of IMC1 results in defects in the lytic cycle

To assess the effects of disrupting *IMC1* on parasite growth, we first performed standard plaque assays (Fig 3A). We found that the Δ*imc1* strain was dramatically hindered in the lytic cycle, displaying a severe 85.7% reduction in plaque area, which was fully rescued by complementation (Fig 3B). We also observed a 71.4% reduction in plaque efficiency, indicating that many of the knockout parasites suffered defects in the lytic cycle that resulted in lethality (Fig 3C). The plaque efficiency defect was again fully rescued in the IMC1^c^ parasites. This extreme decrease in plaque area and efficiency demonstrates that IMC1 is important for one or more stages of the lytic cycle.

**Figure 3.**
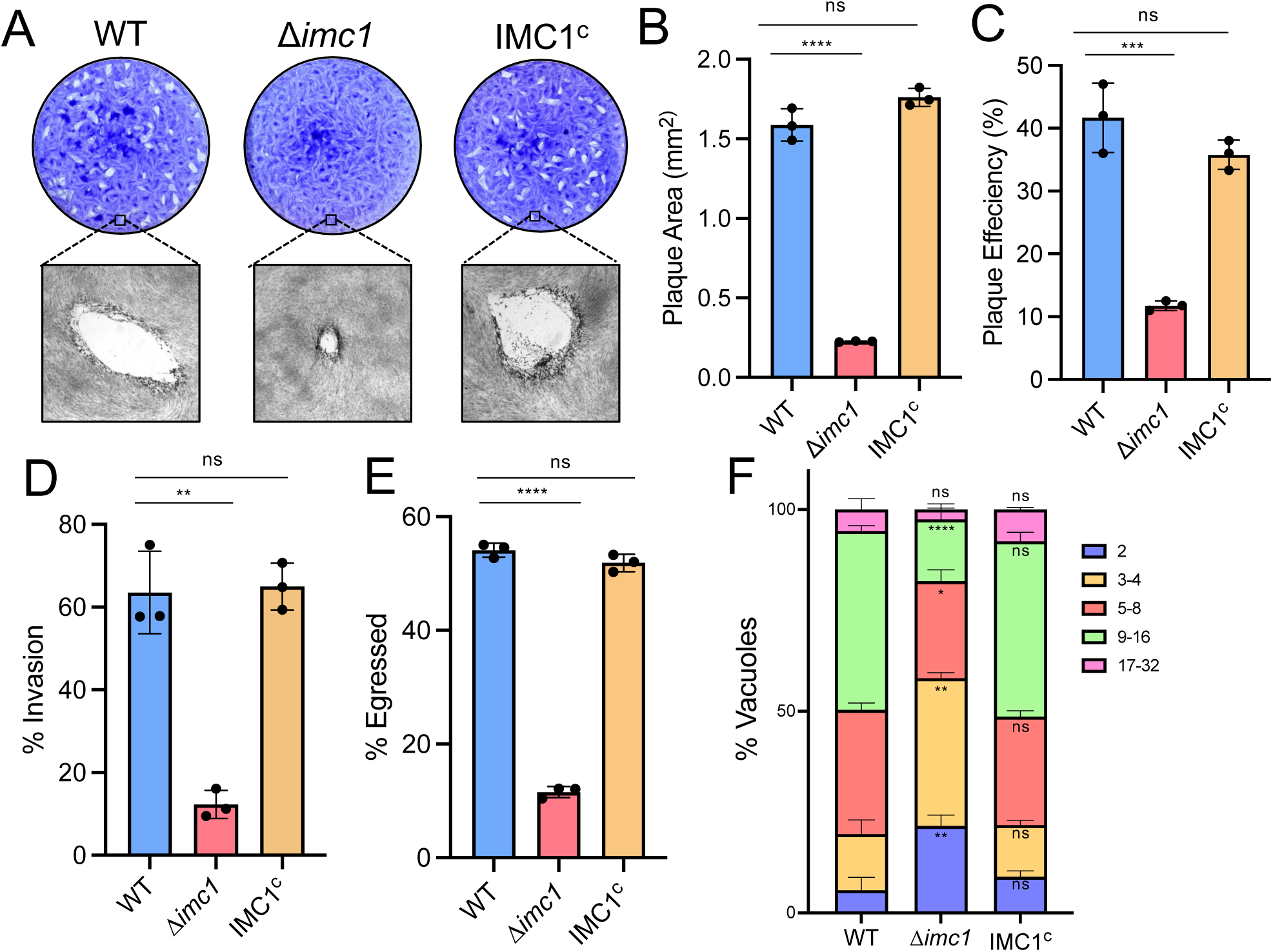
Δ*imc1* parasites have severe defects in the lytic cycle. A) Plaque assays of wild-type, Δ*imc1*, and IMC1^c^ parasites. B) Quantification of plaque area, showing a severe defect in growth in the Δ*imc1* strain. **** P < 0.0001. C) Quantification of plaque efficiency. *** P = 0.0007. D) Quantification of invasion assay, showing % invaded. ** P = 0.0011. E) Quantification of the egress assay, showing percent total vacuoles egressed. **** P < 0.0001. G) Quantification of the replication assay, showing the breakdown of parasites per vacuole for the three strains. **** P < 0.0001. ** P = 0.0026, 0.0018. * P = 0.0229. Statistical significance was determined using two-tailed t tests.

To determine which stages of the lytic cycle were disrupted by the loss of IMC1, we individually examined invasion, replication and egress. We first carried out invasion assays in which we allowed wild-type, Δ*imc1*, and IMC1^c^ parasites to settle on a monolayer in invasion-restrictive media, then altered the conditions to invasion permissive media (31). We then quantified intracellular and extracellular parasites and found that Δ*imc1* parasites showed a severe failure to invade, with only 10% being found intracellular, compared to the wild-type or IMC1^c^ strains, with 63.51% and 64.98% intracellular, respectively (Fig 3D). We also examined the efficiency of parasite egress in the IMC1 knockout using ionophore-induced egress assays (32). We allowed the three strains to form large vacuoles and then induced egress using the calcium ionophore A23187 and observed that 54.1% of the wild-type and 51.85% of the IMC1^c^ vacuoles were induced to egress, while only 11.53% of the Δ*imc1* were (Fig 3E). We then performed replication assays where we allowed the strains to grow for 30 hours (25). We found that the majority of wild-type and complemented parasites had replicated intracellularly to 16 parasites per vacuole, with an average of 44.3% and 43.4% respectively. The Δ*imc1* strain, on the other hand, had only replicated mostly to four-parasite vacuoles each, with an average of 36.6% of total vacuoles being at this stage (Fig 3F). This indicates that the Δ*imc1* strain has defects in replication compared to the wild-type and IMC1^c^ strains. The deficiencies in invasion, egress, and replication indicate that the growth defects caused by the absence of IMC1 are due to a failure to efficiently undergo the lytic cycle at all stages.

### Disruption of IMC1 results in defects in endodyogeny and parasite morphology

To determine exactly how the loss of IMC1 impacts replication, we examined intracellular parasites for defects in morphology and daughter cell formation. We observed multiple defects in endogeny, including asynchronous replication, maternal parasites with more than two daughter buds (multidaughters), disorganized vacuoles, breaks in the IMC network, and incomplete septation (Fig 4A). We quantified the prevalence of these defects in intracellular parasites which strikingly resulted in an average of 99.4% of the Δ*imc1* vacuoles having one or more of these defects while only 2.2% and 2.4% of wild-type and IMC1^c^ vacuoles had any defects, respectively (Fig 4B). We also noticed that, in extracellular parasites, many individual knockout parasites retained the “incomplete septation” defect, with 30.08% of all Δ*imc1* parasites having more than one apical cap, as seen by ISP1 staining, compared to 0.00% and 0.25% of wild-type and IMC1^c^ respectively (Fig 4C, D).

**Figure 4.**
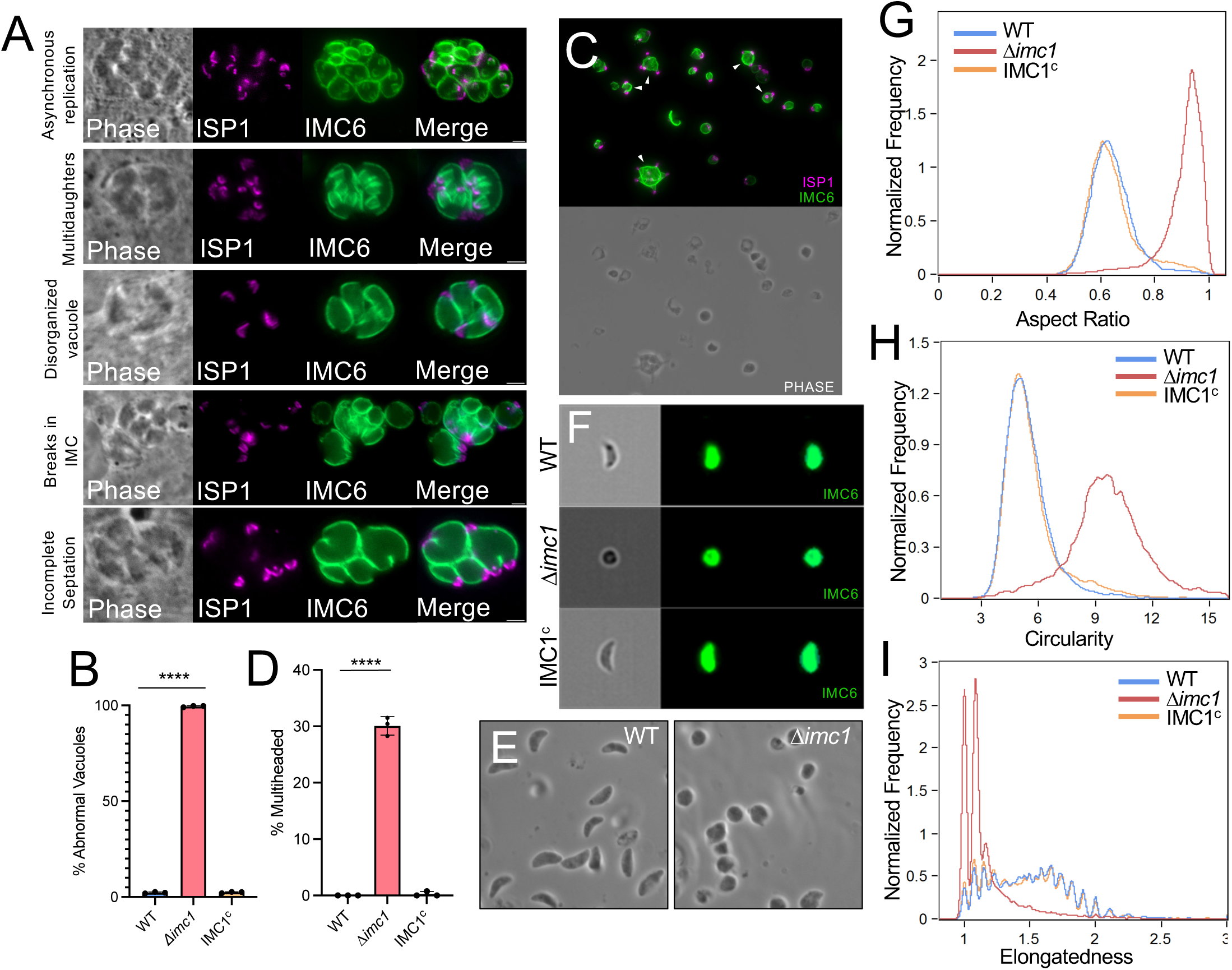
Δ*imc1* displays severe defects in endodyogeny and parasite morphology. A) IFAs of representative images of five different abnormalities in endodyogeny seen in Δ*imc1* parasites stained with mouse anti-ISP1 in magenta and rabbit anti-IMC6 in green. B) Quantification of vacuoles in WT, Δ*imc1*, and IMC1^c^ strains that display at least one or more of the endodyogeny abnormalities shown in Fig 4a. C) IFA of extracellular Δ*imc1* parasites stained with mouse anti-ISP1 in magenta and rabbit anti-IMC6 in green showing multiheaded parasites (arrowheads). D) Quantification of extracellular parasites from WT, Δ*imc1*, and IMC1^c^ strains that are multiheaded. E) Phase contrast image of extracellular wild-type and Δ*imc1* parasites demonstrating extreme rounding in the knockout. F) Representative images captured by ImageStream flow cytometry of WT, Δ*imc1* and IMC1^c^ parasites. Parasite bodies were stained with rabbit anti-IMC6 in green. G) ImageStream analysis of population aspect ratios. Higher aspect ratio values indicate parasite roundness. H) ImageStream analysis of population circularity. Higher circularity values indicate more circular cells. I) ImageStream analysis of population elongatedness. Higher elongatedness values indicate more elongated cells. **** P < 0.0001. Statistical significance was determined using two-tailed t tests.

Examination of extracellular Δ*imc1* parasites by phase contrast microscopy showed severe morphological changes, which was observed as a loss of their normal crescent shape and the formation of strikingly rounded parasites (Fig 4E). To quantify these shape defects, we used ImageStream flow cytometry to image ∼10,000 extracellular parasites of wild-type, Δ*imc1*, and IMC1^c^ that were stained with antibodies to IMC6 to label the periphery of the parasites (Fig 4F) (33). We specifically examined changes in aspect ratio, circularity, and elongatedness and found that Δ*imc1* parasites had an average aspect ratio value of 0.93, compared to 0.63 for both the wild-type and IMC1^c^ strains (Fig 4G). A value closer to 1 suggests a rounder cell, indicating that *Δimc1* displays a significantly rounder morphology. Similarly, Δ*imc1* parasites had an average circularity score of 9.60 while the wild-type and IMC1^c^ strains parasites had scores of 5.24 and 5.26, respectively (Fig 4H). Finally, we observed Δ*imc1* were less elongated than the wild-type and IMC1^c^ strains (Fig 4I). These results show that the loss of IMC1 results in rounder and shorter parasites that have substantial defects in endodyogeny.

### Disruption of IMC1 in type II parasites disrupts the acute and chronic infection in vivo

To determine how the loss of IMC impacts the acute and chronic infection in vivo, we disrupted IMC1 in the Prugnaiud (PruΔ*hxgprt*Δ*ku80*) strain of *T. gondii,* which also contains a *ldh2*-GFP reporter for assessing bradyzoite formation (Fig S2A) (34, 35). The knockout was verified by PCR (Fig S2B) and the resulting Δ*imc1*_II_ strain showed similar morphological defects as the type I knockout (Fig S2A). To verify that Δ*imc1*_II_ parasites retained their ability to form cysts, we performed an in vitro bradyzoite switching assay. The proportion of bradyzoite cyst formation in wild-type and Δ*imc1*_II_ parasites was quantified, which confirmed the knockout strain’s cyst forming ability (Fig 5A). In fact, we found that the Δ*imc1*_II_ strain was more able to switch in vitro with 90.4% of Δ*imc1* vacuoles becoming GFP^+^ while only 39.5% of the wild-type strain were GFP^+^ (Fig 5B). This increase in pH driven switching efficiency may be the result of increased stress already occurring in the knockout strain.

**Figure 5.**
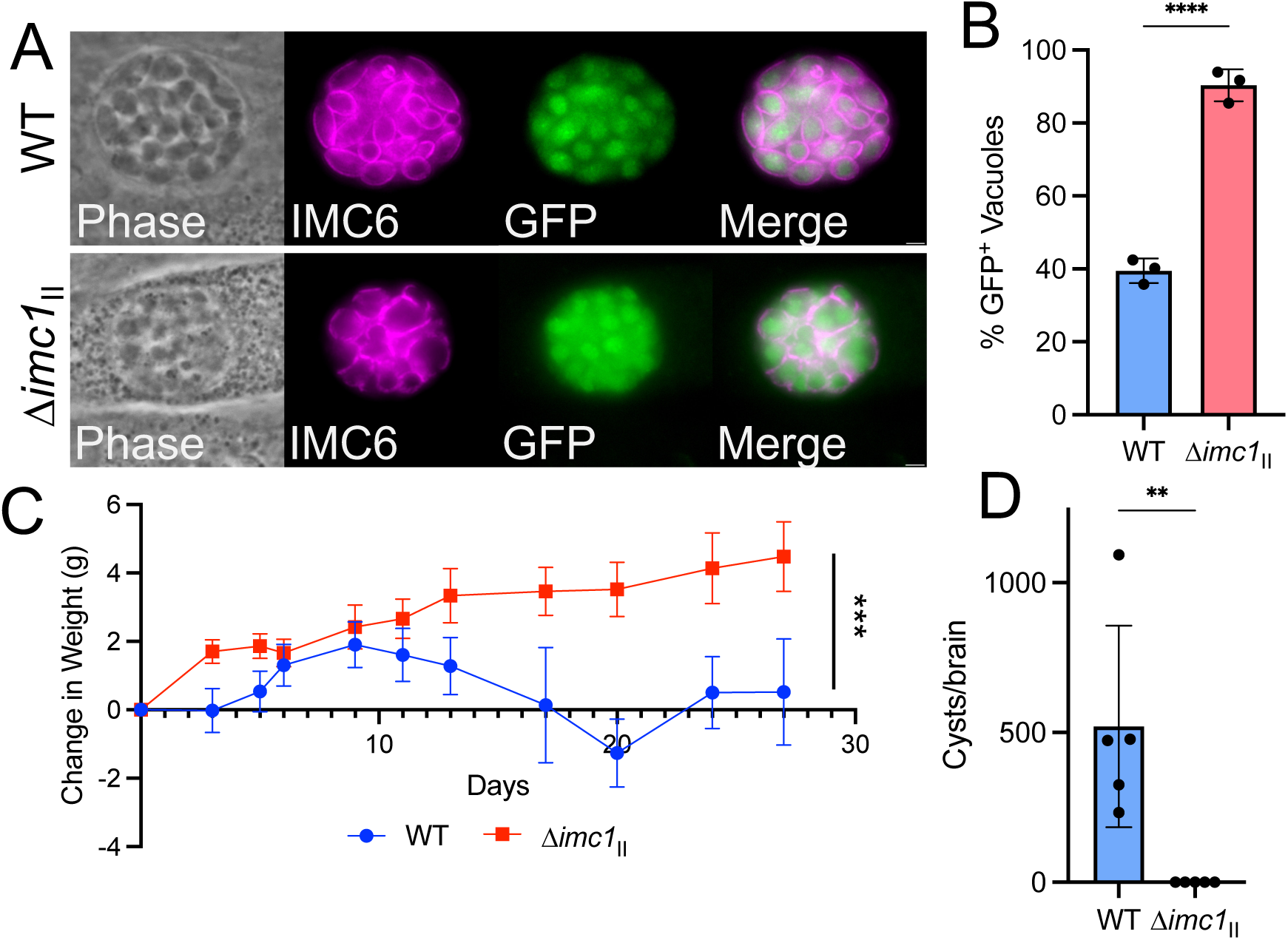
Δ*imc1*_II_ fails to develop a chronic infection *in vivo*. A) IFA of *in vitro* switched GFP^+^ parasite tissue cysts in both WT (top) and Δ*imc1*_II_ (bottom) parasites. B) Quantification of the percentage of GFP^+^ vacuoles after being induced to switch at 93 hpi. C) Time course of average weights of mice infected with WT (Pru*ΔhxgprtΔku80*) parasites (blue circles) and Δ*imc1*_II_ parasites (red squares) through the duration of the infection. E) Quantification of brain cyst burden of mice infected with WT and Δ*imc1* parasites at 30 days post infection. Cyst burden value was normalized for brain weights. Significance was determined using two-tailed t tests. *** P = 0.0003. ** P = 0.0086. **** P < 0.0001. All scale bars are 2 µm. hpi, hours post infection.

To assess the Δ*imc1*_II_ strain’s ability to infect and form cysts in vivo, we infected 5 mice each with an average of 114 and 1235 plaque-forming units of wild-type and Δ*imc1*_II_ parasites, respectively. The mice infected with the wild-type parasites showed a typical weight loss pattern of infection while those infected with the knockout continually gained weight (Fig 5C). After 30 days, the mice were euthanized and serum and brains were collected for serology and cyst enumeration. We found that the mice infected with wild-type parasites resulted in an average of 520 cysts per brain, while no cysts could be found in those infected with Δ*imc1*_II_ parasites (Fig 5D). In addition, we attempted to infect fibroblasts with 50% of the brain homogenate and parasites not could be recovered. We also probed western blots of wild-type parasite lysates with serum from each mouse and found no reactivity from the knockout while sera from the wild-type infection showed a typical signal of infection (Fig S2C, D). To determine if higher doses were better able to establish an infection, we injected three mice with ∼200,000 pfu of the Δ*imc1*_II_ strain and found that the mice were able to seroconvert but again no cysts could be observed or parasites recovered from the brains at 30 days post infection (Fig S2E). Taken together, this indicates that Δ*imc1*_II_ parasites are capable of forming tissue cysts in vitro, but are cleared early in infection and fail to establish a chronic infection in vivo.

### Deletion and mutation analyses elucidates functional domains of IMC1

IMC1 is composed of a conserved alveolin domain in the center of the protein that is flanked by N and C terminal extensions that contain predicted palmitoylation sites (36). Previous work from our lab and others have shown that the alveolin domain is important for proper protein trafficking and function in the alveolins that have been studied (23, 25). To identify regions of IMC1 that are necessary for localization and function, we first assessed the predicted palmitoylation sites at residues C9, C12, and C13 by mutating them from cysteines to alanines in our full-length IMC1 complementation construct (Fig 6A). We then generated an N-terminal deletion series of IMC1, guided by secondary structure and homology to *Plasmodium spp*. (Fig S3A, B) (37). Each of the mutants were targeted to the UPRT locus, as was done for the IMC1^c^ strain. IFA analysis showed that these mutants all localize to the IMC and generally appear to rescue the swollen morphology of the knockout (Fig 6B). Plaque assays showed that the growth defects caused by the knockout are largely rescued by each of the mutants, demonstrating that the N-terminal half of the protein is mostly dispensable (Fig 6C).

**Figure 6.**
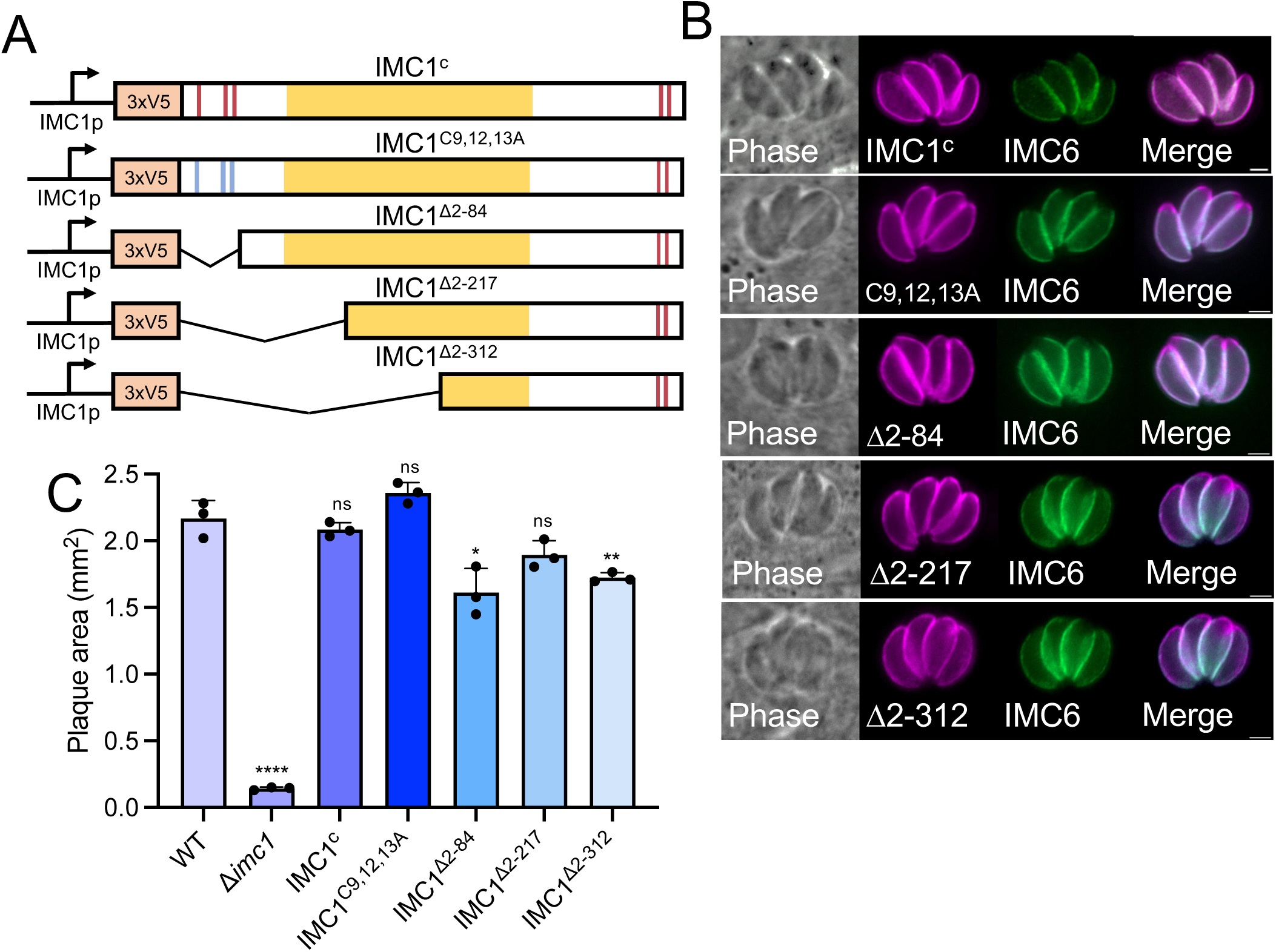
N-terminal mutation and deletion series of IMC1. A) Schematic for mutants created from a full length IMC1 complementation construct. IMC1^C9,12,13A^, IMC1^Δ2-84^, IMC1^Δ2-217^, and IMC1^Δ2-312^. B) IFA of the IMC1 mutants expressed in the Δ*imc1* strain. Each of the constructs targets similar wild-type to the IMC. Magenta is mouse anti-V5, which targets each IMC1 construct, and green is rabbit anti-IMC6. C) Quantification of plaque size for each Δ*imc1* line containing one of the mutant copies of IMC1. IMC1 mutants largely rescue growth defects seen in Δ*imc1* parasites. Statistical significance was determined using two-tailed t tests. **** P < 0.0001. ** P = 0.0063. * P = 0.0131.

Since these mutants removed approximately two-thirds of the alveolin domain, we wanted to determine if we could remove the entire alveolin domain before losing proper IMC1 function (IMC1^Δ2-423^, Fig 7A). We found that deletion of the N-terminal region including the alveolin domain fails to target to the IMC and mislocalizes to the cytoplasm (Fig 7B). We also assessed the growth of these parasites and found that this construct was unable to rescue the Δ*imc1* strain growth defects (Fig 7C). To determine if this region is sufficient for targeting and function, we expressed just this region of the protein in the Δ*imc1* strain and found that this portion localizes properly but the parasites retain the swollen morphology shared by the knockout (IMC1^313-423^, Fig 7D, E). We then generated a mutant that contains all of IMC1 except this region (IMC1^Δ313-423^, Fig 7F) and found that this deletion partially targets to the IMC, but also localizes to the cytoplasm (Fig 7G). When assessing the growth of Δ*imc1* parasites with these mutants, we found that the IMC1^313-423^ strain formed slightly larger plaques compared to the Δ*imc1* strain but still had a sharp defect compared to wild-type parasites (Fig 7H). In contrast, IMC1^Δ313-423^ was unable to rescue the defects of the Δ*imc1* strain. Taken together, these results demonstrate that IMC1^313-423^ is sufficient for IMC targeting and is required for function of the full-length protein.

**Fig 7.**
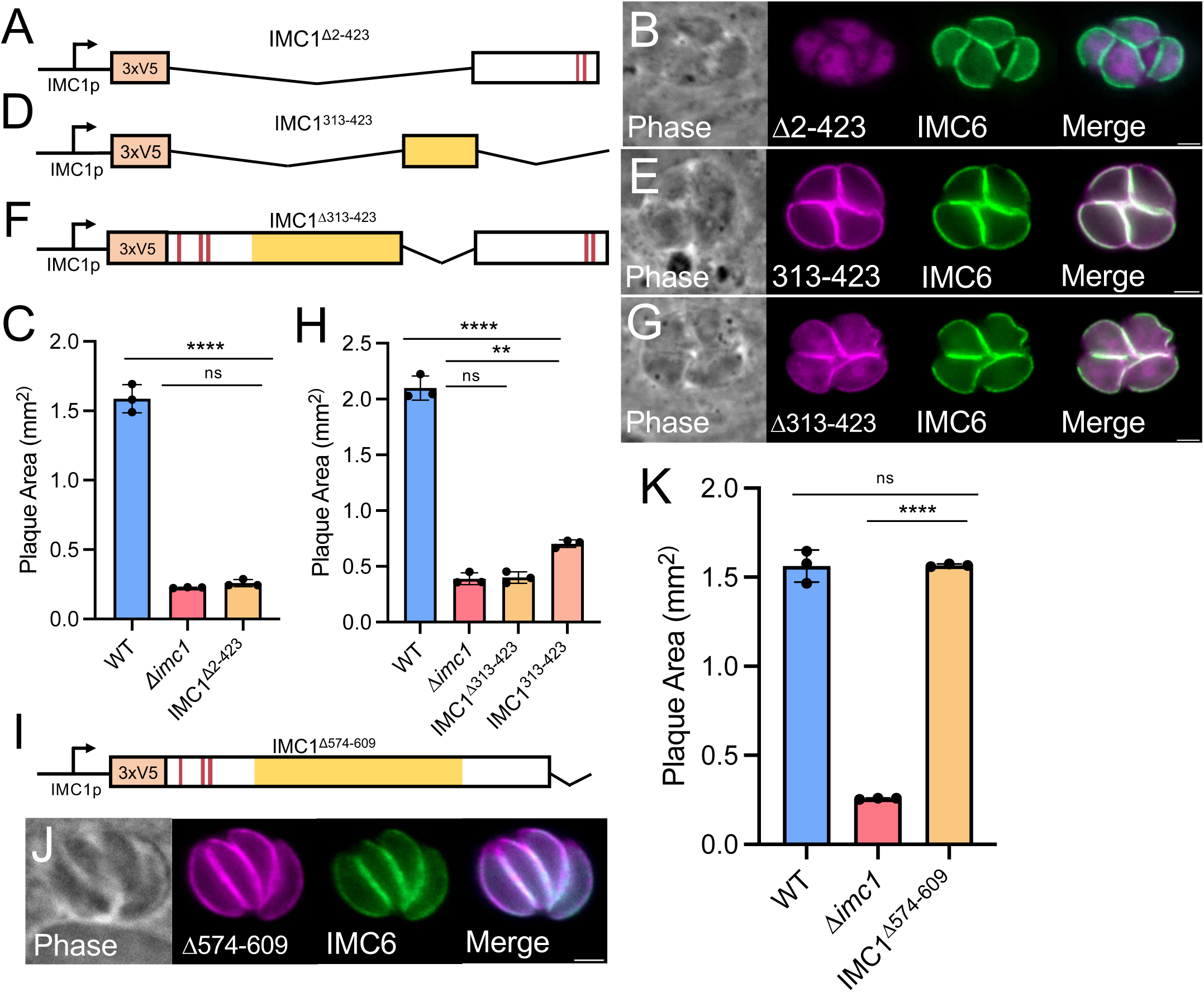
Deletion analyses identify regions of IMC1 important for targeting and function. A) Schematic for the N-terminal deletion removing the entire alveolin domain (IMC1^Δ2-423^). B) IFA showing the IMC1^Δ2-423^ protein is mislocalized to the cytoplasm of the parasite and Δ*imc1* parasite rounding is retained. C) Quantification of plaque area showing the IMC1^Δ2-423^ strain does not rescue growth. D) Schematic for the IMC1^313-423^ mutant which removes all but the C-terminal portion of the alveolin domain. E) IFA of the IMC1^313-423^ mutant within the Δ*imc1* background showing correct localization but parasite rounding retained. F) Schematic of the IMC1^Δ313-423^ mutant that removes the C-terminal portion of the alveolin domain. G) IFA of the IMC1^Δ313-423^ mutant within the Δ*imc1* background showing partial IMC localization and parasite rounding. H) Quantification of the plaque area for the IMC1^Δ313-423^ and IMC1^313-423^ strains. I) Schematic of the IMC1^Δ574-609^ construct. J) IFA of the IMC1^Δ574-609^ strain showing proper localization of IMC1 lacking its C-terminal region and that parasite morphology appears restored. K) Quantification of plaque area showing the IMC1^Δ574-609^ mutant restores the growth defects of Δ*imc1* parasites. Significances were determined using two-tailed t tests. **** P < 0.0001. ** P = 0.0010. All IFA staining is with mouse anti-V5 in magenta and rabbit anti-IMC6 in green.

Previous work has shown that the C-terminal region of IMC1 contains predicted palmitoylation sites and that this region is proteolytically processed, an event which is important for incorporation into the cytoskeletal network of the IMC (29). To determine if this region is important for trafficking and function in Δ*imc1* parasites, we deleted the C-terminal 35 amino acids that encompass the palmitoylation sites from our full-length complementation construct and expressed this deletion mutant in the knockout (Fig 7I). We found that the protein targets to the IMC and mostly rescues the knockout (Fig 7J, K). This data shows that this region is largely dispensable for targeting and function of IMC1.

### Disruption of IMC1 weakens the stability of the parasite cytoskeleton

To determine whether the absence of IMC1 impacts the stability of other alveolins in the cytoskeleton, we performed detergent fractionation experiments to separate the membrane and cytoskeletal elements of the parasite (38, 39). Wild-type and Δ*imc1* strains were individually extracted with 1% Triton X-100, and the solubilized membrane and insoluble cytoskeletal fractions were run on western blots probing for ISP3, a marker of the membrane fraction, and the alveolins IMC3, IMC4, IMC6, and IMC10 for the cytoskeleton. We found that in wild-type parasites, IMC3, IMC4, IMC6, and IMC10 remain in the insoluble pellet while ISP3 is released into the supernatant, as expected (Fig 8A) (39). However, in the Δ*imc1* strain, IMC3, IMC4, IMC6, and IMC10 can be seen in both the supernatant and pellet fractions, suggesting that the lack of IMC1 results in the partial disassociation of these proteins off the cytoskeleton when exposed to detergent (Fig 8B). We next wanted to determine how sensitive the Δ*imc1* parasites were to detergents in general and also compare this to our previous Δ*imc6* strain (25). We thus treated extracellular wild-type, Δ*imc1*, and Δ*imc6* parasites with a solution of 1% Triton X-100 and 1% deoxycholate and stained for tubulin for the cytoskeleton and SAG1 for the plasma membrane. We observed that this treatment resulted in the expected ghosts in both wild-type and Δ*imc6* parasites, in which the cytoskeleton remained relatively intact and SAG1 was released, although the Δ*imc6* strain was misshapen as we previously observed (25). However, using these conditions, Δ*imc1* parasites were clearly more disrupted, often forming umbrella-like structures with the microtubules splayed outwards, suggesting that the cytoskeleton of Δ*imc1* is more sensitive to detergent extraction (Fig 8C). We also examined these strains by transmission electron microscopy and saw similar results with the Δ*imc1* strain being more readily disrupted upon detergent extraction (Fig 8D).

**Figure 8.**
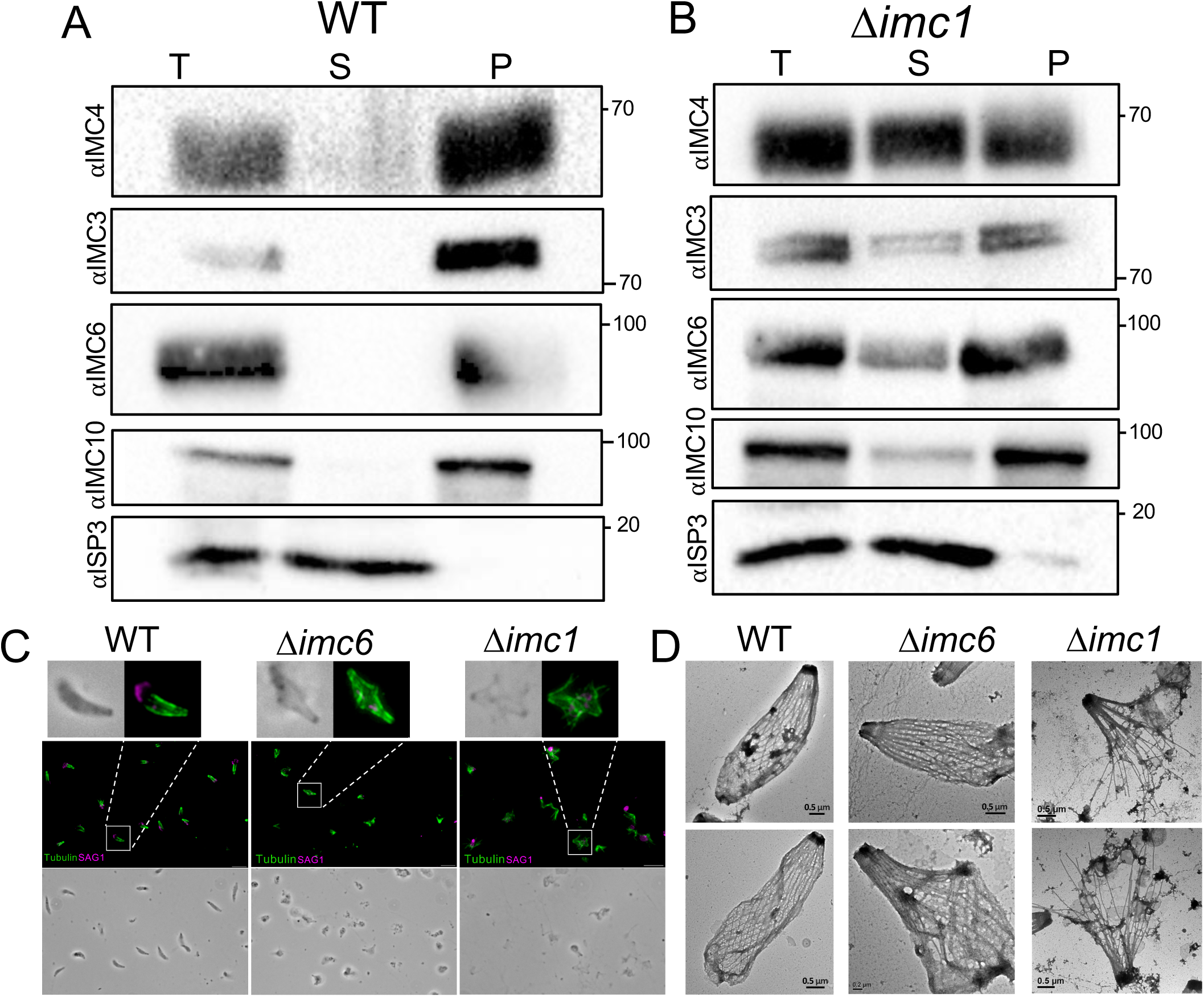
Loss of IMC1 results in heightened cytoskeletal sensitivity to detergent. A-B) Detergent fractionation of A) wild-type and B) Δ*imc1* parasites, demonstrating that IMC3, IMC4, IMC6, and IMC10 are present in both soluble and insoluble fractions of Δ*imc1* parasites treated with 1% Triton X-100. C) IFA of detergent extraction of wild-type (left), Δ*imc6* (middle), Δ*imc1* (right) parasites stained with mouse anti-SAG1 in magenta and rabbit anti-PolyE in green demonstrating the change in detergent sensitivity between Δ*imc1* and Δ*imc6* strains after being treated with 1% Triton X-100 + 1% DOC solution. DOC, deoxycholate. D) TEM images of negative-stained, detergent-extracted wild-type (left), Δ*imc6* (middle), Δ*imc1* (right) parasites showing the Δimc1 strain SPN is more disrupted by detergent extraction.

### IMC4 is mislocalized in Δ*imc1* parasites and IMC1 and IMC4 directly interact

Given the morphological changes and increased sensitivity to detergent extraction in Δ*imc1* parasites, we next examined whether a series of other alveolins (IMC3, IMC4, IMC6, IMC7, IMC10, and IMC12) were potentially impacted upon loss of IMC1. While IMC3, IMC6, IMC7, IMC10 and IMC12 localize normally to the IMC (Fig S4B-D), IMC4 – which normally overlaps perfectly with IMC1 – is mislocalized exclusively in the forming daughter buds of replicating Δ*imc1* parasites (Fig 9A, B). We then tested this with the IMC1^Δ313-423^ and IMC1^313-423^ complemented strains as well and found that, while IMC4 is mislocalized in the IMC1^Δ313-423^ strain (Fig 9C), its localization is restored to wildtype in the IMC1^313-423^ parasites (Fig 9D). This indicates that residues 313-423 of IMC1 play an important role in IMC4 recruitment during daughter bud formation. Since IMC1 and IMC4 co-localize with each other in wild-type parasites, this defect in the absence of IMC1 demonstrates a coordination of IMC1 and IMC4 within the parasite cytoskeleton.

**Figure 9.**
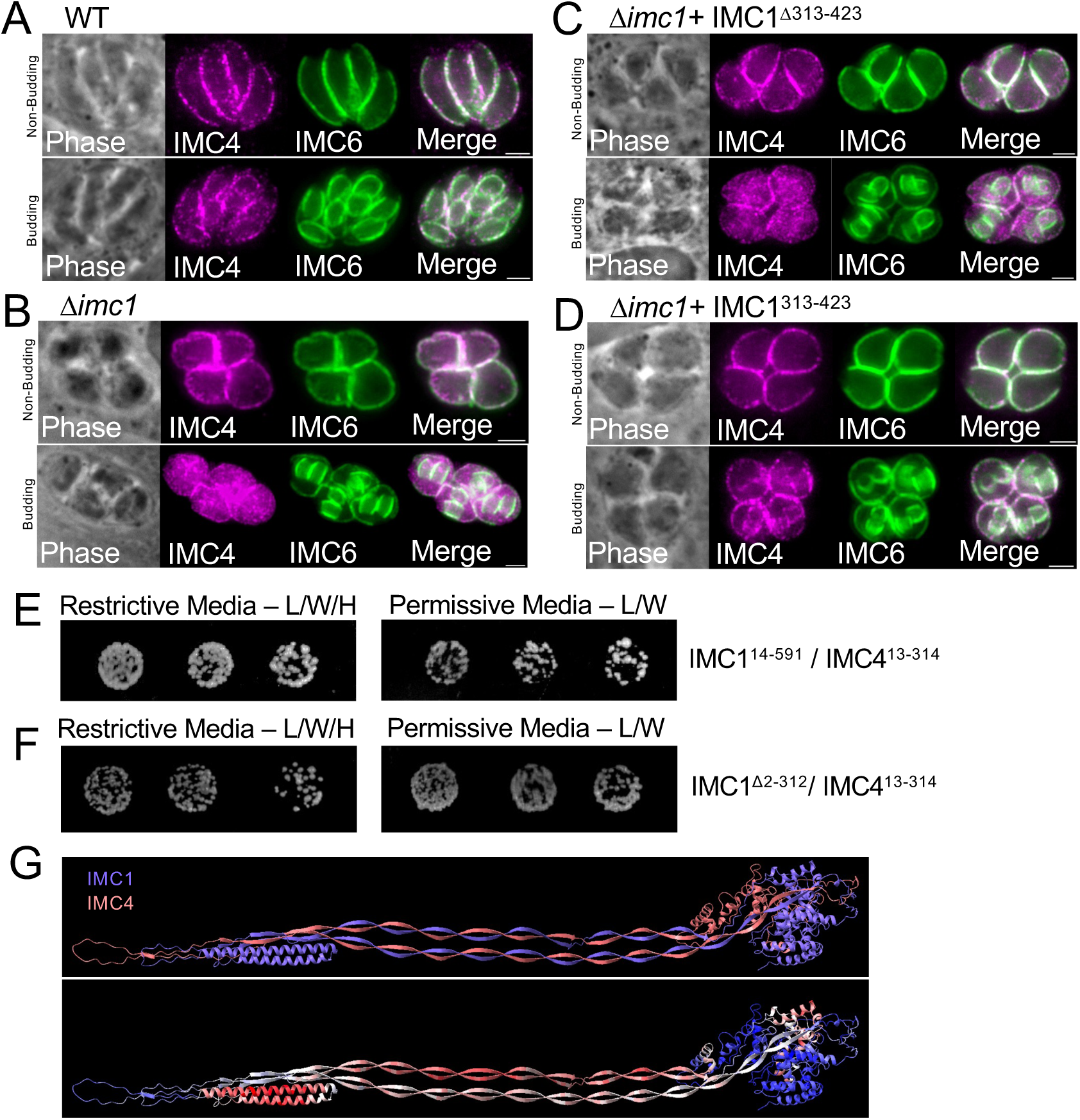
IMC4 is mislocalized in Δ*imc1* daughter buds and directly binds IMC1. A) IFA of non-budding (top) and budding (bottom) wild-type parasites stained with rat anti-IMC4 and rabbit anti-IMC6 showing proper localization of IMC4. B) IFA of Δ*imc1* parasites stained with rat anti-IMC4 and rabbit anti-IMC6 showing mislocalization of IMC4 in daughter buds. C) IFA of IMC1^Δ313-423^ showing IMC4 remains mislocalized in daughter buds. D) IFA of IMC1^313-423^ demonstrating that residues 313-423 are sufficient for the association of IMC4 to the growing daughter bud IMC. E, F) Images of yeast strain growth on restrictive (left) and permissive media (right) demonstrating IMC1 and IMC4 interact. G) Proposed binding interaction between IMC1 and IMC4 (top) and confidence prediction (bottom) as predicted by AlphaFold. Red indicates higher confidence, blue indicates lower confidence, b-factor ranges from 20.8 to 92.3.

Based on these results, we wanted to determine if IMC1 binds directly to IMC4. To address this question, we performed pairwise yeast two-hybrid assays. To do this, the IMC1 coding region lacking the regions containing its predicted palmitoylation sites (IMC1^14-591^) was cloned into the pP6 (GAL4 activation domain) prey vector and residues 13-314 of IMC4 were cloned into the pB27 (LexA) bait vector (16). The constructs were co-transformed into yeast and growth was assessed on either permissive or restrictive media. We found that the IMC1/IMC4 transformed yeast grew on restrictive media, indicating a direct interaction (Fig 9E). To determine whether the smallest of our N-terminal deletion series that functions can bind IMC4 (see Fig 6A, IMC1^Δ2-312^), we cloned this portion into the bait vector and found that it could also interact with IMC4 (Fig 9F). Autoactivation was tested and confirmed that none of these constructs autoactivated when paired against the corresponding empty vector (Fig S4E). Together, this data demonstrates that IMC1 interacts directly with IMC4 and this interaction is present in the C-terminal half of the protein, likely in the alveolin domain. To better understand how IMC1 and IMC4 may interact to form filament-like structures, we used AlphaFold to predict their potential organization (40). This suggested that the beta sheets of the alveolin domain interact with one another, and the proteins are organized in a parallel fashion when forming the IMC network filaments (Fig 9F). In addition, the interwoven alveolin domains of IMC1 and IMC4 appear to fold in upon each other and form electrostatic interactions between beta sheets of the alveolin domains.

## Discussion

In this work, we investigate the function of IMC1, the founding member of the *Toxoplasma* alveolins that form the SPN (28). Like most IMC proteins, the alveolins are expressed in a “just in time” fashion during endodyogeny in which components are sequentially expressed and loaded onto the forming daughter buds (13). Our analysis of the expression timing of IMC1 and IMC4 shows that these alveolins overlap perfectly, but appear significantly later than IMC3, IMC6 and IMC10. The earlier expression and enrichment of IMC3, IMC6 and IMC10 in daughter buds suggests that these are important for the establishment of the IMC network to support the IMC32/IMC43/BCC0 “essential daughter assembly complex” along with other early daughter expressed IMC proteins (14–16). The increased intensity of IMC3, IMC6 and IMC10 in early buds may suggest that most of the material that is needed in maternal parasites is expressed at this early stage, which then becomes more less prominent as it diffuses into the larger maternal IMC. The IMC1 and IMC4 pair are then added later, suggesting they function in IMC network assembly in the later stages of daughter IMC development or are primarily important in maternal parasites.

Despite IMC1’s strongly negative phenotype score, we were able to disrupt it, demonstrating that it is not strictly essential for parasite survival (26). However, loss of IMC1 results in severe morphological changes to the parasite that has dramatic effects on endodyogeny and also impacts invasion and egress. We previously demonstrated similar morphological effects upon disruption of the daughter-enriched alveolin IMC6, but the Δ*imc*1 parasites appear even more rounded and less pointed in the apical cap area of the IMC (25). This is likely due to IMC1 and IMC4 extending into the apical cap and providing structure in this location, whereas IMC3, IMC6 and IMC10 are all restricted to the body of the IMC. Another key difference is that Δ*imc1* parasites have an egress defect while Δ*imc6* parasites do not, suggesting that maternal IMC functions are partially disrupted. Interestingly, there are two orthologues of IMC1 in *Plasmodium spp*. named IMC1a and IMC1b, which are expressed in sporozoites and ookinetes, respectively (41, 42). Similar to our findings, disruption of IMC1a results in shape, tensile strength and infectivity defects in sporozoites and disruption of IMC1b results in morphological defects and reduced mechanical strength, motility and infectivity of ookinetes. It is likely that redundancy exists for these proteins in other lifecycle stages in *Plasmodium*.

We also use Δ*imc1*_II_ parasites to explore how these defects impact infectivity and tissue tropism in vivo and find that moderate doses of the knockout do not cause illness and are likely cleared as the infected mice do not seroconvert and cannot establish a chronic infection (35). While high doses of the knockout do result in seroconversion, the parasites cannot be detected in the brain in the chronic infection, indicating these also are likely cleared during the earlier stages of infection. It would be interesting to disrupt *IMC6* in a type II background to determine if these are also cleared or can persist longer in vivo.

Our deletion and mutagenesis experiments reveal regions of IMC1 that are necessary for IMC targeting and function. The N-terminal and C-terminal palmitoylation sites are largely dispensable for IMC1 localization and function, suggesting that other alveolins or other SPN associated proteins contain IMC membrane anchors that are sufficient for tethering the SPN to the IMC membranes. It is also possible that either group of these sites is sufficient for membrane tethering or that these sites play a minor role that is not detected in our assays. The C-terminal region also includes the proteolytic processing event that mediates maturation of IMC1, which has been proposed to be important for incorporation into the maternal IMC network (29). Our C-terminal deletion that removes this region questions its importance, but it is possible that our deletion mimics this cleavage event.

Surprisingly, the N-terminal half of the protein, including ∼2/3 of the alveolin domain is also largely dispensable for both targeting and function. To our knowledge, this is the first functional dissection of an alveolin domain in *T. gondii*. The IMC1 alveolin domain from *Toxoplasma* shares substantial sequence homology with its *Plasmodium* orthologues IMC1a and IMC1b in the N-terminal and C-terminal regions of the domain with a noticeable gap in sequence similarity between these regions (Fig S3). Interestingly, sequence analyses of the *Plasmodium* alveolin domains identified two conserved subdomains (termed type I or type II) within IMC1a and IMC1b that correlate with these regions of homology (43). Our deletion series demonstrates that the N-terminal region of homology corresponding to the type I domain is largely dispensable, while the C-terminal region containing the type II domain contains the region that we have shown is both sufficient for localization to the IMC and necessary for function.

We also show that loss of IMC1 leads to a weakened ultrastructure of the IMC in that Δ*imc1* parasites are more sensitive to detergent extraction as assessed by IFA and electron microscopy. This more fragile network is also observed via the partial release of several alveolins upon detergent extraction. The partial destabilization of the network is distinct from that seen in Δ*imc6* parasites which share similar morphological and replication defects but are less prone to dissociation in detergent (25). These results indicate that IMC1 likely plays a more important role in stability of the maternal SPN in extracellular parasites. Interestingly, loss of IMC1 also results in a mistargeting of IMC4 in forming daughter buds by IFA. This, along with the perfect colocalization of IMC1 and IMC4, suggested that these alveolins are partners, which we verified by pairwise yeast two-hybrid analysis. We also attempted to determine if IMC4 could be disrupted but were unable to obtain knockout parasites, indicating that it is essential. This agrees with recent studies in *Plasmodium*, which show that the IMC4 orthologue IMC1g is essential for parasite viability (44, 45). While IMC4 is mislocalized in forming daughter buds of Δ*imc1* parasites, it properly localizes to the maternal IMC, suggesting that it likely interacts with other components of the maternal cytoskeleton.

Together, our results and those of others suggests that there are two critical complexes of the key alveolins that are present during replication. The first group to be expressed are IMC3, IMC6, IMC10, and ILP1, which all are enriched in daughter buds and appear prior to IMC1 and IMC4. We have previously shown that IMC3, IMC6 and ILP1 interact, and IMC10 is likely to be a member of this complex based on its overlapping expression pattern. The second group is IMC1 and IMC4, which are expressed in later daughters and show similar expression levels in daughter buds and maternal parasites. It is likely that the two groups interact in late daughters and/or maternal parasites given the extensive interactions of the alveolins described so far. We also attempted to disrupt IMC6 in the Δi*mc1* strain, but were unable to obtain double knockouts, suggesting that the combined effects of these knockouts are too disruptive to the IMC network and results in lethality. Future work will focus on the precise architecture of these proteins, how they are regulated, and how their disassembly and degradation is governed during each round of replication.

## Materials and Methods

### T. gondii culture

Parental *T. gondii* RHΔ*hxgprt*Δ*ku80* (46), Pru*Δku80Δhxgprt (ldh2GFP)* (35), and subsequent strains were cultured on confluent monolayers of human foreskin fibroblasts (HFF) host cells at 37°C and 5% CO_2_ in Dulbecco’s Modified Eagle Medium (DMEM) supplemented with 5% fetal bovine serum (Gibco), 5% Cosmic Calf serum (HyClone), and 1x penicillin-streptomycin-l-glutamine (Gibco). Strains were selected using media supplemented with either 50 µg/mL of mycophenolic acid/xanthine (MX) (47) or 5µg/mL of 5-Fluoro-5’-deoxyuridine (FUDR) (30).

### Gene Knockout

To generate a knockout of IMC1, a protospacer was designed to target an exon within IMC1 (TgGT1_231640) and ligated it into the pU6-Universal plasmid utilizing primers P1-2 (Table I) (48). The homology-directed repair (HDR) template included 40 bp of homology immediately upstream of the start codon and 40 bp of homology approximately 200bp downstream of the stop codon (P3-4). Along with an HXGPRT selectable marker driven by a NcGRA7 promoter, the HDR template was amplified from a pJET vector using P3-4. For transfection, the PCR-amplified HDR template was purified by phenol-chloroform extraction and this plus ∼50 µg of the gRNA-carrying pU6-Universal plasmid were ethanol precipitated. Both constructs were electroporated into the RHΔ*hxgprt*Δ*ku80* or *PruΔku80Δhxgprt (ldh2GFP)* parental parasite strains. Transfected parasites were allowed to invade a confluent monolayer of HFFs overnight, and appropriate selection was subsequently applied. Successful knockout was confirmed by IFA, and clonal lines were obtained through limiting dilution.

### IMC1 Complementation

All IMC1 complementation constructs were modified from the previously generated IMC29 complementation construct (49). Each construct contains either the full-length or truncated version of the IMC1 coding sequence and UPRT homology regions to drive this cassette to the *UPRT* locus. For this study, the IMC29 promoter was replaced with the IMC1 promoter using Gibson assembly. The IMC1 promoter was amplified from genomic DNA (P13-14). The full length IMC1 cDNA and the vector plasmid were amplified using P9-12. The product was purified and ligated using the NEBuilder HiFi DNA Assembly kit, resulting in the final plasmid. The full-length construct was also remade with an additional N-terminal 3xV5 epitope tag. To make truncation constructs, the same process was followed for each truncation using the tagged version: IMC1^Δ2-84^ was amplified using P22/P21, IMC1^Δ2-217^ was amplified using P23/P21, IMC1^Δ2-312^ was amplified using P24/P21, IMC1^Δ2-423^ was amplified using P25/P21, and IMC1^Δ574-609^ was amplified using P27/P26 off of the full-length constructs. Additionally, IMC1^313-423^ was amplified using P28/P26 from IMC1^Δ2-312^ and IMC1^Δ313-423^ was amplified using P29/P30 from the full-length plasmid. After plasmid construction, 100 µg of each complementation plasmid was linearized with PsiI-HF and transfected into Δ*imc1* parasites with a pU6-gRNA that targets the UPRT coding region. Selection was performed with 5ug/ml FUDR for replacement of *UPRT* and clones were screened by IFA (30).

### Antibodies

The V5 epitope tag was detected using mouse mAb anti-V5 (catalog no. R96025; Invitrogen). *T. gondii*-specific antibodies used include mouse mAb anti-IMC1 (50), pAb rabbit anti-IMC3 (51), rabbit pAb anti-IMC6 (22), rat pAb anti-IMC6 (22), rat pAb anti-IMC7 (24), rat pAb anti-IMC10 (27), rabbit pAb anti-IMC12 (52), mAb mouse anti-SAG1 DG52 (53), mouse mAb anti-MIC2 (54), and rabbit pAb anti-SAG2 (55), mouse mAb anti-ROP7 (56), mouse mAb anti-ISP1 (46), mouse mAb anti-F_1_β subunit (57), rabbit pAb anti-CENTRIN1 (Kerafast; EBC004), rat pAb DrpB (58), mouse mAb anti-Atrx1 (59), mouse pAb anti-ISP3 (46), and rabbit pAb anti-catalase (60).

### Immunofluorescence assay and western blot

For immunofluorescence assays, HFF cells were seeded on glass coverslips and allowed to grow until confluency. Parasites were then allowed to infect these monolayers and 18– 36 hours post infection, the coverslips were fixed with 3.7% formaldehyde in PBS and processed for immunofluorescence as described (60). Primary antibodies are added to blocking buffer (1x PBS, 5% BSA, 0.2% TX-100) and incubated on the coverslips for one hour, washed with PBS five times for two minutes each, species-specific secondary antibodies (Alexa Fluor 488/594) were incubated for one hour, and washed with PBS 5 times. Coverslips were then mounted on a microscope slide in VectaShield (Vector Labs, Burlingame, CA), viewed with an Axio Imager.M2 fluorescent microscope (Zeiss), and processed with Zeiss ZEN 3.7 software (Zeiss), which included deconvolution. Images with the 594-fluorophore channel were pseudocolored to magenta.

For western blotting, parasites were lysed using 1x Laemmli Sample Buffer (50 mM Tris-HCL [pH 6.8], 10% glycerol, 2% SDS, 100 mM DTT, and 0.1% bromophenol blue) and boiled for 10 minutes. Lysates were run on SDS-PAGE gels and transferred to nitrocellulose membranes. The membranes were blocked using 5% milk and 1xPBS supplemented with 0.1% Tween-20 mixture for half an hour before probing with the appropriate primary antibody and the corresponding secondary antibody conjugated to horse radish peroxidase (HRP). Probing was visualized on ChemiDoc XRS+ (Bio-Rad) using SuperSignal chemiluminescence substrate (Thermo Scientific).

### Detergent fractionation

To separate parasites into detergent soluble and insoluble fractions, parasites were collected, washed with 1xPBS, and resuspended in 1% Triton X-100 supplemented with Complete Protease Inhibitor Cocktail (Roche) (38, 39). This was incubated on ice for 20 minutes before being centrifuged at 14,000g for 15 minutes to separate the sample. The supernatant was mixed with 2x Laemmli Sample Buffer and the pellet was resuspended in an equivalent volume of 1x Sample Buffer. The fractions were then visualized using SDS-PAGE and Western Blotting. Antibodies probing for IMC6 were used to detect the insoluble cytoskeletal fraction and ISP3 was used to indicate the solubilized membrane fraction.

### Detergent extraction of extracellular parasites

For extracellular detergent extraction the parasites were washed in 1xPBS were first allowed to settle on coverslips coated in poly-L-lysine for 30 minutes. The remaining liquid was then aspirated and replaced with 1% Triton X-100 and 1% DOC and allowed to incubate for 30 minutes. The coverslips were then washed and fixed with 3.7% formaldehyde in 1xPBS for 15 minutes. The coverslips were then washed again, stained with primary and secondary antibodies, and visualized by fluorescence microscopy. For transmission electron microscopy, parasite ghosts were prepared using an on-grid extraction technique for wild-type, Δ*imc1* and Δ*imc6* parasites (25). Briefly, 4 μl of parasite resuspension was applied onto a glow-discharged continuous carbon-coated EM grid. After allowing the sample to settle for 2 minutes, the excess buffer was blotted from the edge of the grid. The grid was then placed face-down onto a 20 μl drop of detergent extraction buffer (1% Triton X-100, 0.5% DOC, 50 mM Tris, 150 mM NaCl) and left to float for 5 minutes to lyse the parasites. After this initial extraction, the grid was blotted again and transferred to a fresh drop of the same buffer for another 5 minutes. This extraction step was repeated six times in total. Subsequently, the grid was washed four times using 20 μl drops of PBS, following the same procedure. Finally, the grid was stained with 2% uranyl acetate for negative staining and imaged using a FEI Tecnai T12 transmission electron microscope equipped with a CCD camera and operated at 120 kV.

### Plaque assay

Six-well plates hosting HFF monolayers were allowed to become confluent. Then, equivalent number of parasites were allowed to infect the monolayer for seven days, forming plaques. The samples were fixed with ice-cold methanol and stained with crystal violet (24). For each condition, the area of around 50 plaques were measured using ZEN blue software. Additionally, the number of plaques formed by the parasites added was quantified. All plaque assays were performed with three biological replicates. Statistical analysis, graphs, and figures were generated using Prism GraphPad 10.

### In Vitro Bradyzoite Switching Assay

Parasites were allowed to invade 24-well plates with confluent HFFs for three hours before changing media into a high pH media designed to induce *in vitro* switching (61). The coverslip plate was then allowed to sit at 5% CO_2_ and 37°C for three hours. The plate was then wrapped in parafilm to seal any remaining CO_2_ inside of the wells and moved to an incubator at 0% CO_2_ and 37°C for 12 hours. After 12 hours, the plate was returned to an incubator at 5% CO_2_ and 37°C for one hour before repeating the process until 93 hours had elapsed. The coverslips were then probed with anti-IMC6 antibodies and the experiment was quantified by determining the percentage of vacuoles that had turned into cysts (GFP^+^) out of the total vacuoles in each field. Triplicate experiments were performed, with each replicate spanning at least 15 fields and 50 vacuoles. Statistical analysis, graphs, and figures were generated using Prism GraphPad 10.

### Mouse Infections and Brain Cyst Quantification

Wild-type PruΔ*ku80*Δ*hxgprt* and Δ*imc1* parasites were counted and resuspended in Opti-MEM prior to intraperitoneal injection into groups of five female CBA/J mice each. A total of 114 wild-type and 1235 Δ*imc1* plaque forming units/mouse were used for the innoculum.

For high dose infections, ∼200,000 pfu were used of the Δ*imc1* strain for infection. The mice were observed for 30 days and then sacrificed. Mouse brains were collected, homogenized, and examined for the presence of LDH2-GFP+ tissue cysts (34). Quantification was performed by examining 25-µl aliquots of each brain homogenate using fluorescence microscopy until approximately 25% of the brain by volume was counted. Total cyst burden was then extrapolated. Statistical analysis, graphs, and figures were generated using Prism GraphPad 10.

### Replication Defect Quantification

HFFs were seeded on glass coverslips until fully confluent and were infected at low MOI, incubated for 3 hours at 37°C, and extracellular parasites were washed away by media changes. Then, 30 hours post-infection, coverslips were fixed with 3.7% formaldehyde, processed for immunofluorescence, and labeled with anti-ISP1 and anti-IMC6. To score normal vs. abnormal vacuoles, >300 vacuoles across at least 15 fields were counted. Vacuoles were categorized as abnormal if any one of the replication defects shown in Fig 4 were present in a vacuole. Statistical analysis, graphs, and figures were generated using Prism GraphPad 10.

### Invasion assay

Parasites in large vacuoles were mechanically lysed using a syringe and resuspended in invasion restrictive media and settled onto coverslips with a HFF monolayer for 20 minutes (31). The invasion restrictive media was then replaced with warm growth media supplemented with 20 mM HEPES and incubated in 37°C and 5% CO_2_ for 30 minutes.

Coverslips were then fixed, blocked with non-permeabilizing 3% BSA in 1x PBS for 30 minutes. The extracellular parasites were stained with anti-SAG1 antibodies for an hour. The coverslips were washed with 1xPBS and permeabilized with 3% BSA, 0.2% Triton X-100, 1x PBS for 30 minutes, and all parasites were stained with anti-SAG2. Coverslips were washed with 1xPBS and mounted on microscope slides using the mounting medium VectaShield (Vector Labs) and viewed with an Axio Imager.M2 fluorescence microscope (Zeiss). Parasites were assigned with SAG1-,SAG2+ (Invaded) or SAG1+,SAG2+ (Uninvaded). Three biological replicates were performed with at least 300 parasites across 15 fields per triplicate. Statistical analysis, graphs, and figures were generated using Prism GraphPad 10.

### Egress Assay

HFFs were infected at a low MOI and grown until vacuoles reached 16-32 parasites per vacuole. The media was then changed to warm HBSS –/+ 1uM A23187 ionophore (32). Fixative was added to the coverslips after 4 minutes and the coverslips were stained with the rabbit anti-IMC12. Percentage of egressed vacuoles per total vacuoles was enumerated in triplicates with at least 100 vacuoles and 15 fields per replicate. Statistical analysis, graphs, and figures were generated using Prism GraphPad 10.

### Pairwise Yeast 2-Hybrid Assay

Using Gibson Assembly, genes of interest were cloned into pB27 (N-LexA-bait-C fusion) and pP6 (N-GAL4^AD^-prey-C fusion) vectors containing either a N-terminal GAL4 Activating Domain or a LexA DNA Binding Domain. Smaller fragments were built using Q5 site-directed mutagenesis using primers (P33-41). Pairs of constructs containing both a pB27 and a pP6 were co-transformed into the L40 strain of *S. cerevisiae* [MATa his3D200trp1-901 leu2-3112 ade2 LYS2::(4lexAop-HIS3) URA3::(8lexAop-lacZ) GAL4]. Stains were allowed to grow overnight in liquid permissive media (–Leu/-Trp), re-diluted to OD_600_=2 and then spotted in serial dilutions on permissive (–Leu/-Trp) and restrictive (– Leu/-Trp/-His) plates. After growing for 3-5 days, the plates were imaged. Auto-activation was tested for by co-transforming individual constructs with their corresponding empty vector into yeast and performing spot assays as described previously (16).

### Replication defects by IFA

Parasites were allowed to invade HFFs seeded on a coverslip at a low MOI for 1 hour at 37°C, then remaining extracellular parasites were washed away. Coverslips were then fixed using 3.7% formaldehyde and stained with mouse anti-ISP1 and rabbit anti-IMC6. To score normal vs. abnormal vacuoles, >300 vacuoles across at least 15 fields were counted. Vacuoles were categorized as abnormal if one or more of the replication defects shown in Fig 4A were present in a vacuole. All IFAs were performed in triplicate. For extracellular parasites, fully lysed wild-type, Δ*imc1*, and IMC1^c^ strains were collected and washed in 1xPBS before being settled onto coverslips coated with poly-L-lysine (Sigma Aldrich). These coverslips were then stained using IFA with mouse anti-ISP1 and rabbit anti-IMC6, taking caution not to disturb the parasites. Quantifications were performed by counting three replicates of >100 parasites per strain. Statistical analysis, graphs, and figures were generated using Prism GraphPad 10.

### ImageStream Flow Cytometry

Mechanically lysed extracellular parasites were stained using indirect immunofluorescence in solution. The parasites were fixed using 3.7% formaldehyde, permeabilized and blocked in solution. These parasites were then stained with rabbit anti-IMC6 and anti-rabbit Alexa Fluor 588. For each strain, around 10,000 individual images were taken by the INSPIRE application on the ImageStream MKII (Cytek Biosciences, Seattle, WA) flow cytometer. Images were acquired using a 6 μm width core and 60x magnification. Analysis of the samples were performed using the IDEAs v6.4 software (Cytek Biosciences, California). The aspect ratio of each cell was calculated using the ratio of the minor axis divided by the major axis. The circularity of each cell was measured by a cell’s degree of deviation from a circle, with higher values indicating higher circularity and lower values indicating a higher degree of variation from a circle. Elongatedness was calculated using the ratio of height over width of a cell, with lower values indicating less elongatedness.

### Animal experimentation ethics statement

Our protocol was approved by the UCLA Institutional Animal Care and Use Committee (Chancellor’s Animal Research Committee protocol: #2004–005). Mice were euthanized when the animals reached a moribund state and euthanasia was performed following AVMA guidelines.

## Author contributions

JNU, QL, ZHZ, and PJB designed research; JNU and QL performed the research; JNU, QL, ZHZ, and PJB analyzed data; and JNU, QL, and PJB wrote the manuscript.

## Classification

Biological Sciences – Microbiology

## Acknowledgements

We thank Salem Haile and Zoran Galic of the UCLA Flow Cytometry Core for help acquiring ImageStream data. We thank Vern Carruthers for the MIC2 antibodies, Gary Ward for the IMC1 antibody, Marc-Jan Gubbels for the IMC3 and IMC7 antibodies, Dominique Soldati-Favre for catalase antibodies, and Lloyd Kasper and John Boothroyd for the SAG1 antibody. This work was supported by the NIH grant AI064616 to P.J.B. JNU was supported by the UCLA Undergraduate Scholars Research Program.

## Figure Legends

**Figure S1.** The apicoplast, mitochondria, centrosomes, micronemes, rhoptries, and golgi are unaffected in Δ*imc1* parasites. (A-F) IFAs of wild-type and Δ*imc1* parasites, showing normal morphology of the indicated organelles. (A) The apicoplast was detected with mouse anti-ATrx1 (magenta). (B) Mitochondria were detected with mouse anti-F_1_β (magenta). (C) The centrosome was detected with mouse anti-CENTRIN1 (magenta). (D) Micronemes were detected with mouse anti-MIC2 (magenta). (E) Rhoptries were detected with mouse anti-ROP7 (magenta). (F) The Golgi (trans-Golgi) was detected with rat anti-DrpB (magenta). All IFAs were costained with rabbit anti-IMC6 (green). All scale bars are 2 μm.

**Figure S2.** Gene knockout of *IMC1* in the PruΔ*hxgprtΔku80* strain and infected mice serology. A) IFA of intracellular WT parasites showing proper localization of IMC1 (top). IFA of intracellular Δ*imc1*_II_ parasites showing absence of IMC1 and swollen morphology (bottom, arrows). B) PCR verification of the genomic loci from WT (PruΔ*hxgprtΔku80*) and Δ*imc1*_II_ parasites. Diagram demonstrates the primers used to amplify regions of the IMC1 coding sequence (blue arrows, coding region check) and the regions containing the 5’ site of recombination for the knockout locus (5’ recombination check, red arrows). C, D) Western blots of whole parasite lysates probed with infection sera at 30 days post infection showing that the mice fail to seroconvert with a ten-fold increase (114 vs. 1235 pfu) in infectious dose of the knockout compared to wild-type mice which all seroconvert. E) Seroconversion is obtained at a high dose (∼200,000 pfu) of the Δ*imc1*_II_ strain.

**Figure S3.** Clustal Omega alignment of *Toxoplasma* IMC1 with IMC1a and IMC1b from *Plasmodium falciparum*. A, B) Alignment of *Toxoplasma* IMC1 (TgGT1_231640) with IMC1a (panel A, PF3D7_0304000) or IMC1b (panel B, PF3D7_1141900) from *Plasmodium falciparum*. For *T. gondii*, the predicted palmitoylation sites are highlighted in green, the alveolin domain is highlighted in yellow, and the region that is essential for IMC targeting is highlighted with an orange box. Asterisks indicate identity, a colon indicates conservation between groups of strongly similar properties, and a period indicates conservation between groups of weakly similar properties.

**Figure S4.** Alveolins unaffected in Δ*imc1* parasites and yeast two-hybrid controls. A-D) IFAs showing that IMC3, IMC7, IMC10, and IMC12 are unaffected in both budding and non-budding Δ*imc1* parasites. The proteins are detected with their respective antibodies and IMC6, also unaffected, is used for costaining. E) Controls for yeast two-hybrid experiments with empty vector partners showing a lack of autoactivation by failure to grow on restrictive media (L/W/H). Growth on permissive media (L/W) is also shown.

