## Supplementary Figs 1-4 for "*Toxoplasma* IMC1 is a central component of the subpellicular network and plays critical roles in parasite morphology, replication, and infectivity"

S1 Fig

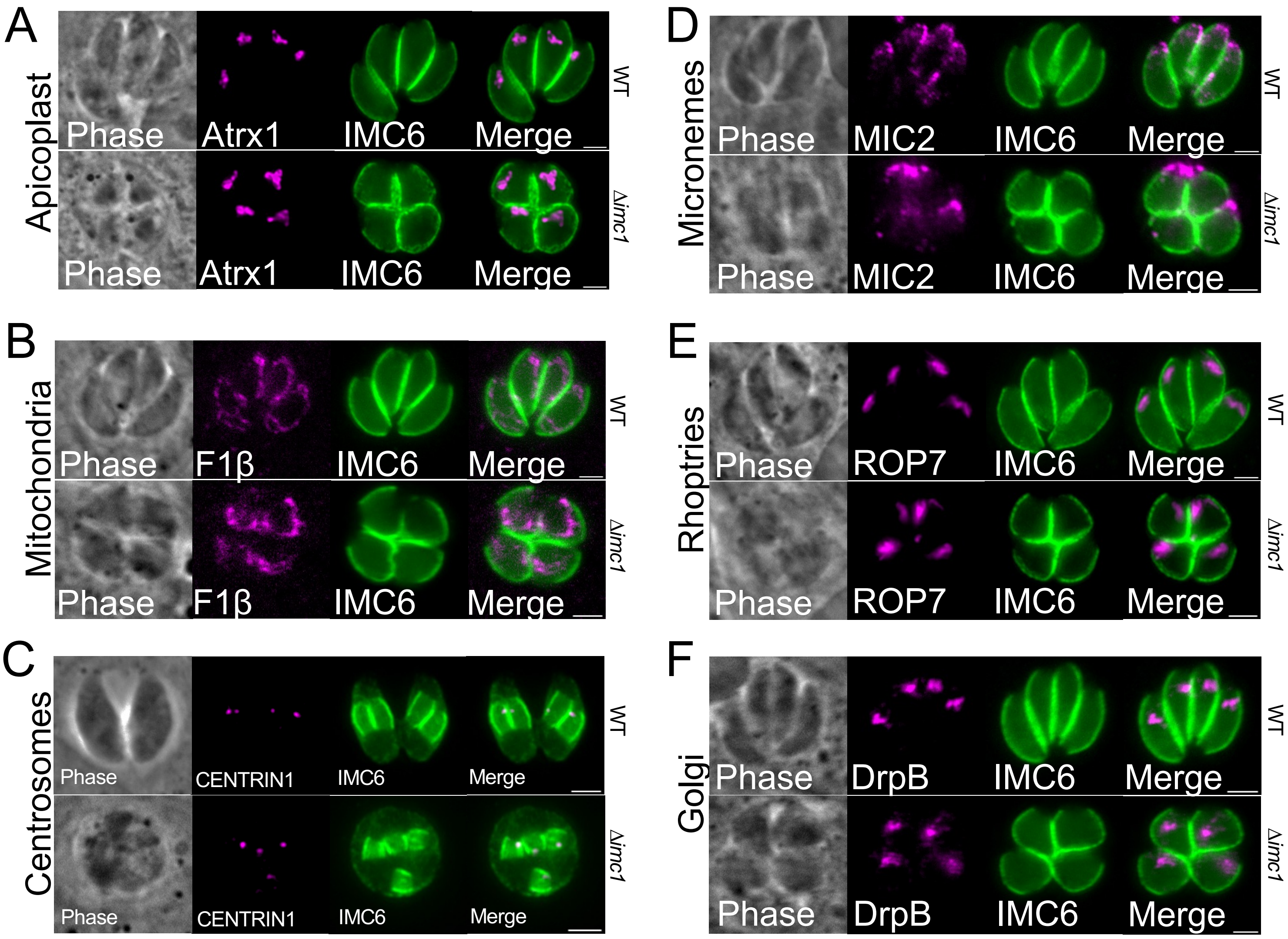


**Figure S1. The apicoplast, mitochondria, centrosomes, micronemes, rhoptries, and golgi are unaffected in Δ*imc1* parasites.** (A-F) IFAs of wild-type and Δ*imc1* parasites, showing normal morphology of the indicated organelles. (A) The apicoplast was detected with mouse anti-ATrx1 (magenta). (B) Mitochondria were detected with mouse anti-$F_1\beta$ (magenta). (C) The centrosome was detected with mouse anti-CENTRIN1 (magenta). (D) Micronemes were detected with mouse anti-MIC2 (magenta). (E) Rhoptries were detected with mouse anti-ROP7 (magenta). (F) The Golgi (trans-Golgi) was detected with rat anti-DrpB (magenta). All IFAs were costained with rabbit anti-IMC6 (green). All scale bars are 2 μm.

### S2 Fig.

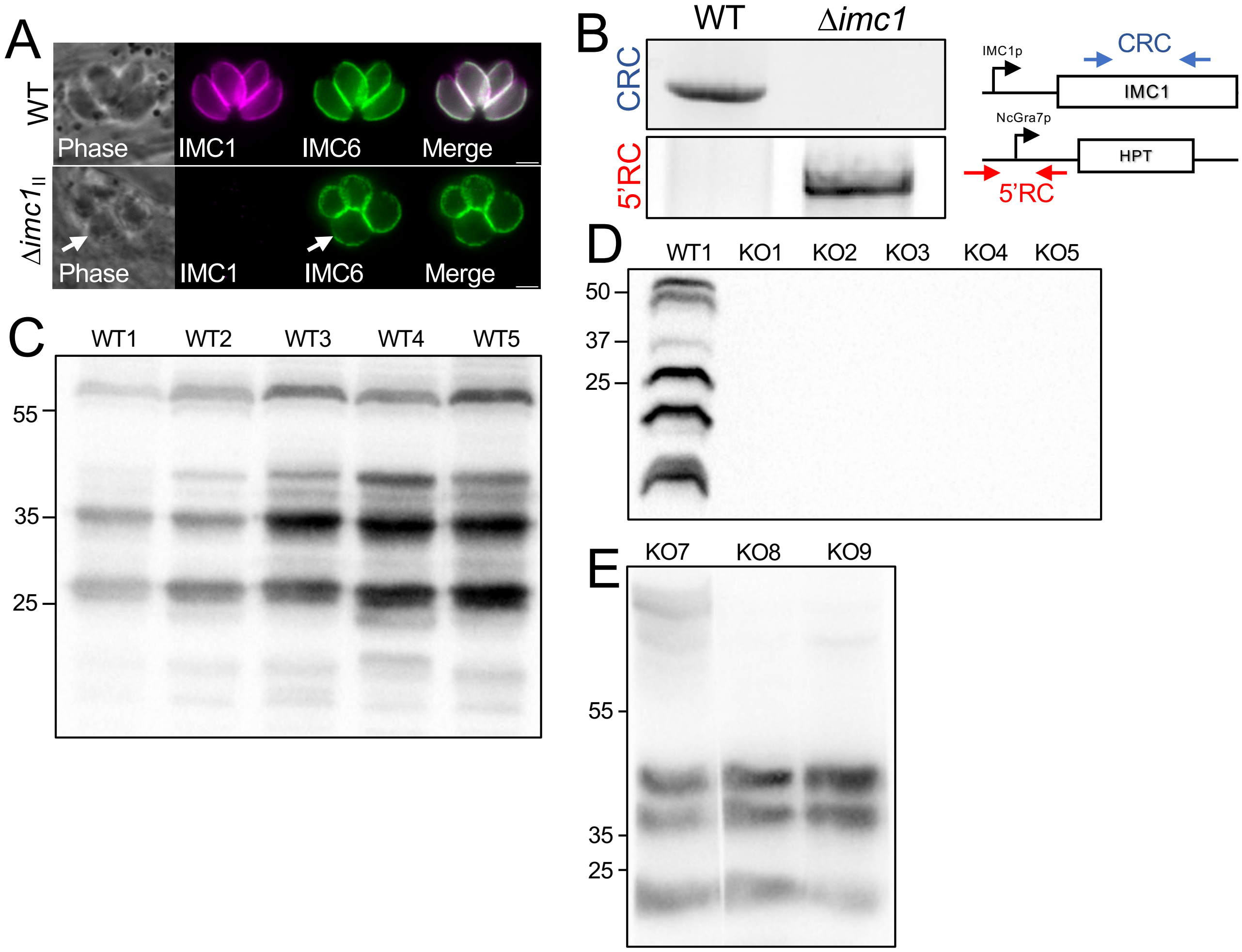


**Figure S2. Gene knockout of *IMC1* in the PruΔ*hxgprt*Δ*ku80* strain and infected mice serology.** A) IFA of intracellular WT parasites showing proper localization of IMC1 (top). IFA of intracellular Δ*imc1*$_{II}$ parasites showing absence of IMC1 and swollen morphology (bottom, arrows). B) PCR verification of the genomic loci from WT (PruΔ*hxgprt*Δ*ku80*) and Δ*imc1*$_{II}$ parasites. Diagram demonstrates the primers used to amplify regions of the IMC1 coding sequence (blue arrows, coding region check) and the regions containing the 5' site of recombination for the knockout locus (5' recombination check, red arrows). C, D) Western blots of whole parasite lysates probed with infection sera at 30 days post infection showing that the mice fail to seroconvert with a ten-fold increase (114 vs. 1235 pfu) in infectious dose of the knockout compared to wild-type mice which all seroconvert. E) Seroconversion is obtained at a high dose (~200,000 pfu) of the Δ*imc1*$_{II}$ strain.

### S3 Fig.

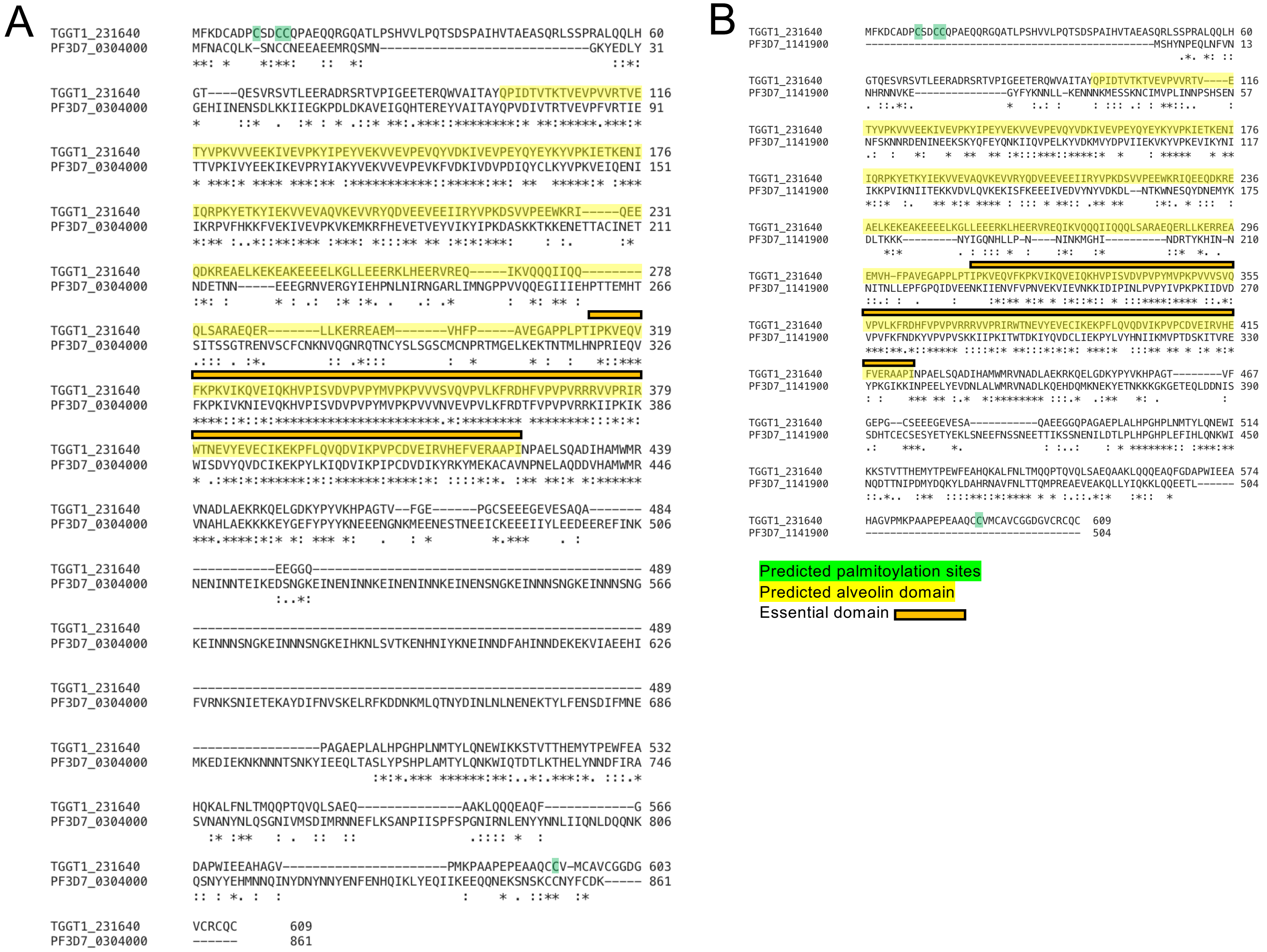


**Figure S3. Clustal Omega alignment of *Toxoplasma* IMC1 with IMC1a and IMC1b from *Plasmodium falciparum*.** A, B) Alignment of *Toxoplasma* IMC1 (TgGT1_231640) with IMC1a (panel A, PF3D7_0304000) or IMC1b (panel B, PF3D7_1141900) from *Plasmodium falciparum*. For *T. gondii*, the predicted palmitoylation sites are highlighted in green, the alveolin domain is highlighted in yellow, and the region that is essential for IMC targeting is highlighted with an orange box. Asterisks indicate identity, a colon indicates conservation between groups of strongly similar properties, and a period indicates conservation between groups of weakly similar properties.

S4 Fig.

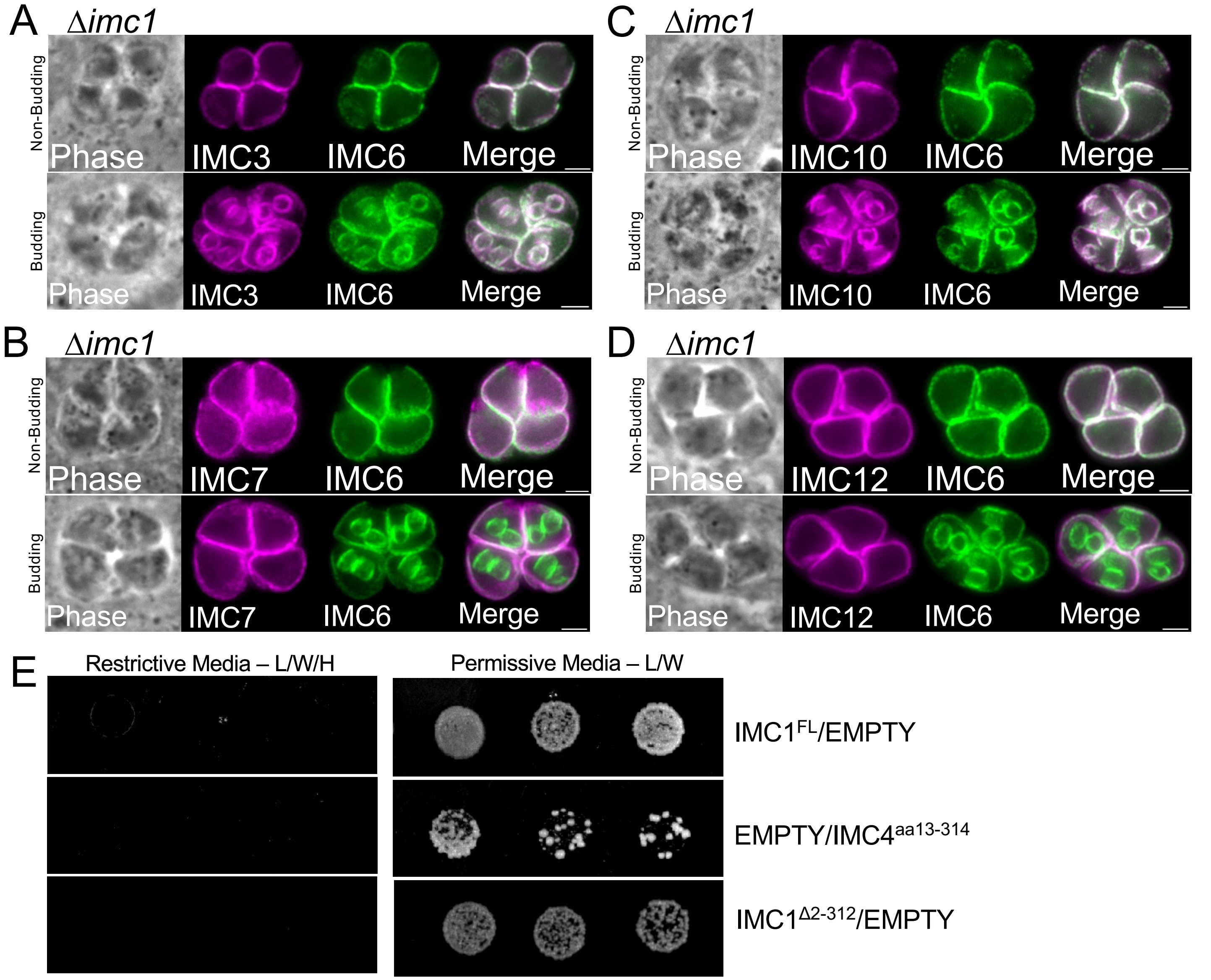


**Figure S4. Alveolins unaffected in Δ*imc1* parasites and yeast two-hybrid controls.** A-D) IFAs showing that IMC3, IMC7, IMC10, and IMC12 are unaffected in both budding and non-budding Δ*imc1* parasites. The proteins are detected with their respective antibodies and IMC6, also unaffected, is used for costaining. E) Controls for yeast two-hybrid experiments with empty vector partners showing a lack of autoactivation by failure to grow on restrictive media (L/W/H). Growth on permissive media (L/W) is also shown.
